# Interpersonal alignment of neural evidence accumulation to social exchange of confidence

**DOI:** 10.1101/2022.11.08.515654

**Authors:** Jamal Esmaily, Sajjad Zabbah, Bahador Bahrami, Reza Ebrahimpour

## Abstract

Private, subjective beliefs about uncertainty have been found to have idiosyncratic computational and neural substrates yet, humans share such beliefs seamlessly and cooperate successfully. Bringing together decision making under uncertainty and interpersonal alignment in communication, in a discovery plus pre-registered replication design, we examined the neuro-computational basis of the relationship between privately-held and socially-shared uncertainty. Examining confidence-speed-accuracy trade-off in uncertainty-ridden perceptual decisions under social vs isolated context, we found that shared (i.e., reported confidence) and subjective (inferred from pupillometry) uncertainty dynamically followed social information. An attractor neural network model incorporating social information as top-down additive input captured the observed behaviour and spontaneously demonstrated the emergence of social alignment in virtual dyadic simulations. Electroencephalography showed that social exchange of confidence modulated the neural signature of perceptual evidence accumulation in the central parietal cortex and produced a sustained top-down flow of information from prefrontal to parietal cortex. Our findings offer a neural population model for interpersonal alignment of shared beliefs.

## Introduction

We communicate our confidence to others to share our beliefs about uncertainty with them. However, numerous studies have shown that even the same verbal or numerical expression of confidence can have very different meanings for different people in terms of the underlying uncertainty[1]–[3]. Similar inter-individual diversity has been found at the neural level [3]–[5]. Still, people manage to cooperate successfully in decision making under uncertainty[6]. What computational and neuronal mechanisms enable people to converge to a *shared meaning* of their confidence expressions in interactive decision making despite the extensively documented neural and cognitive diversity? This question drives at the heart of recent efforts to understand the neurobiology of how people adapt their communication to their beliefs about their interaction partner[7], [8]. A number of studies have provided compelling empirical evidence of brain-to-brain coupling that could underlie adaptive communication of shared beliefs [9]–[15]. These works remain, to date, mostly observational in nature. Plausible neuro-computational mechanism(s) accounting for how interpersonal alignment of beliefs may arise from the firing patterns of decision-related neural populations in the human brain are still lacking [16], [17]. Using a multidisciplinary approach, we addressed this question at behavioral, computational and neurobiological levels.

By sharing their confidence with others, joint decision makers can surpass their respective individual performance by reducing uncertainty through interaction[6], [18]. Recent works showed that during dyadic decision making, interacting partners adjust to one-another by matching their own average confidence to that of their partner[19]. Such confidence matching turns out to be a good strategy for maximizing joint accuracy under a range of naturalistic conditions e.g., uncertainty about the partner’s reliability. However, at present there is no link connecting these socially observed emergent characteristics of confidence sharing with the elaborate frameworks that shape our understanding of confidence in decision making under uncertainty[2], [3], [20]–[26].

Theoretical work has shown that sequential sampling can, in principle, provide an optimal strategy for making the best of whatever uncertain, noisy evidence is available to the agent. These models have had great success in explaining the relationship between decision reaction time and accuracy under a variety of conditions ranging from perceptual[27]–[29] to value-based decisions[30], [31] guiding the search for the neuronal mechanisms of evidence accumulation to boundary in rodent and primate brains[32]. The relation between reaction time and accuracy, known as speed-accuracy tradeoff, has been recently extended to a three-way relationship in which choice confidence is guided by *both* reaction time and probability (or frequency) of correct decision[22], [28], [33]. Critically, these studies have all focused on decision making in *isolated individuals* deciding privately[17]. Little is known about how these computational principles and neuronal mechanisms can give rise to socially shared beliefs about uncertainty.

To bridge this gap, we examined confidence-speed-accuracy tradeoff in Social vs Isolated context in humans. We combined a canonical paradigm (i.e. dynamic random dot motion) extensively employed in psychophysical and neuroscientific studies of speed-accuracy-confidence tradeoff[27], [29], [34], [35] with interactive dyadic social decision making[6], [19]. We replicated the emergence of confidence matching and obtained pupillometry evidence for shared subjective beliefs in our social implementation of the random dot paradigm and we observed a novel pattern of confidence-speed-accuracy tradeoff under specifically under the social condition. We constructed a neural attractor model that captured this tradeoff, reproduced confidence matching in virtual social simulations and made neural predictions about the coupling between neuronal evidence accumulation and social information exchange that were born out by the empirical data.

## Results

We used a discovery-and-replication design to investigate the computational and neuro-biological substrates of confidence matching in two separate steps: 12 participants (4 female) were recruited in Study 1 (Discovery) and 15 (5 female, age: 28 (mean) +/-Std (7)) in study 2 (Replication, second study was pre-registered: https://osf.io/5zces). In each study, participants reported the direction of a random-dot motion stimulus and indicated their confidence (Figure 1a) while EEG and eye tracking data were recorded, simultaneously. After an extensive training procedure (see Methods for the recruitment), participants reached a stable behavioral (accuracy and RT) performance level. Then, two experimental sessions were conducted: first a Private session (200 trials) in which participants performed the task alone; then a Social session (800 trials for study 1 and 400 for study 2) in which they performed the task interactively together with a partner (implied to be another participant in a neighboring lab room).

**Figure 1.**
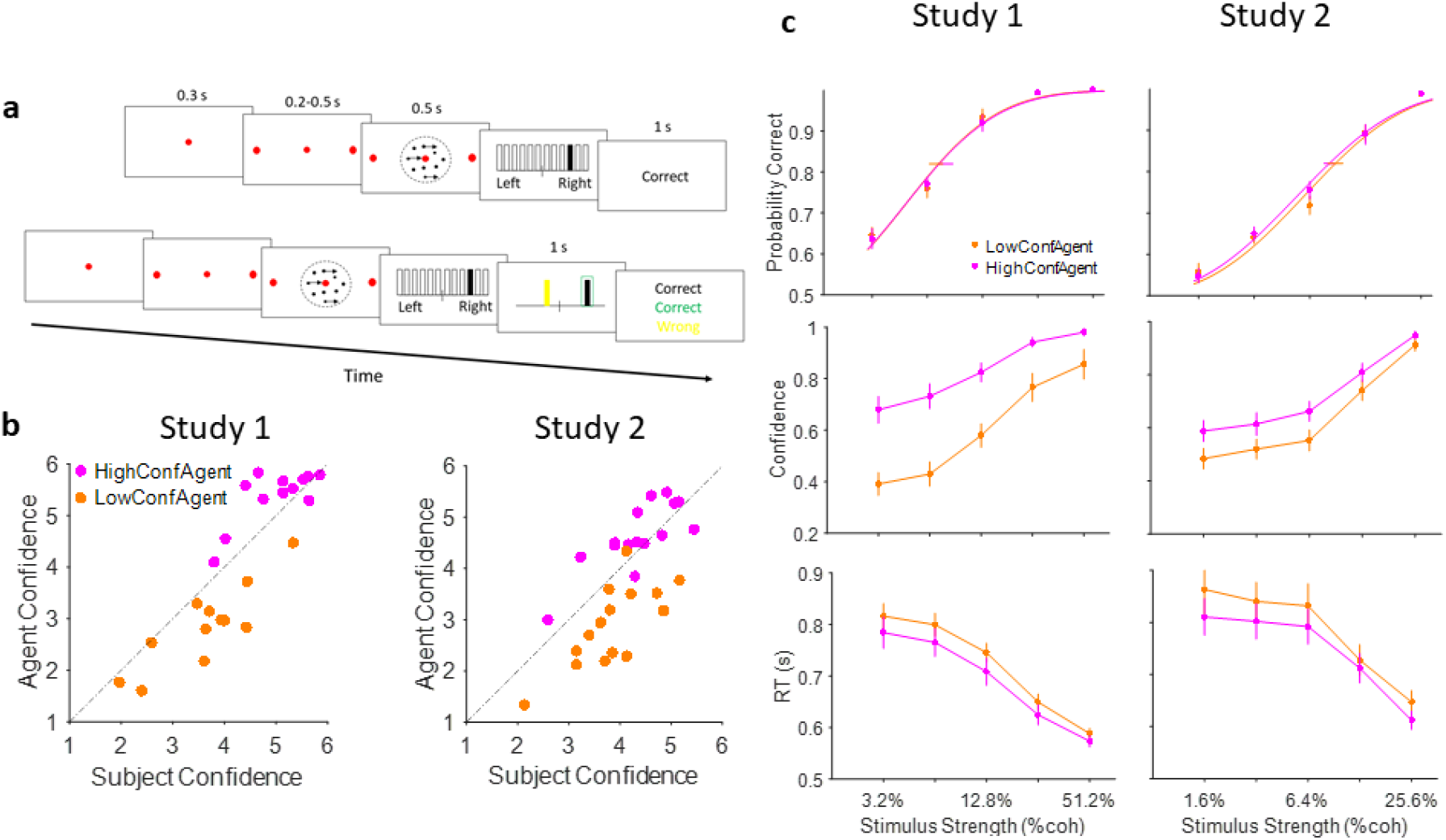
Experiment paradigm and behavioral results. (a) Timeline of trials in Isolated (top) and Social (bottom) conditions. After stimulus presentation, subjects reported their decision and confidence simultaneously by clicking on one of the 12 vertical bars. In the Social condition, decision and confidence of participant (white in the experiment, here black for illustration purpose) and partner (yellow) were color coded. (b) confidence matching. Participants confidence against agent confidence show a significant relation in both studies (linear regression p<0.001 for both studies). (c) Under Social condition, when participants were paired with high (magenta) vs low (dark orange) confidence partner, accuracy (top panel) did not change (horizontal lines, 68 % confidence interval of bootstrap test with 10,000 repetitions) but confidence (middle panel) and RT (bottom panel) were altered. Curves fitted to the accuracy data are Weibull cumulative distribution function. Error bars are standard error of the mean (SEM) across subjects.

In every trial (Figure 1a), after fixation for 300ms was confirmed by closed-loop real-time eye tracking, two choice-target points appeared at 10° eccentricity corresponding to the two possible motion directions (left and right). After a short random delay (200-500ms, truncated exponential distribution), a dynamic random dot motion [see ref [35]] was centrally displayed for 500ms in a virtual aperture (5° diameter). At the end of the motion sequence, the participant indicated the direction of motion and their confidence on a 6-point scale by a single mouse click. A horizontal line intersected at midpoint and marked by twelve rectangles (six on each side) was displayed. Participants moved the mouse pointer – initially set at the midpoint – to indicate their decision (left vs right of midpoint) and confidence by clicking inside one of the rectangles. Further distance from the midpoint indicated more confidence. Reaction time was calculated as the time between the onset of the motion stimulus sequence and the onset of deviation of the mouse pointer (see methods for more details) [36] at the end of stimulus presentation.

In the Isolated trials, the participant was then given visual feedback for accuracy (Correct or Wrong). In the Social trials (Figure 1a bottom panel), after the response, participants proceeded to the social stage. Here, the participants’ own choice and confidence as well as that of their partner were displayed coded by different colors (White for participants; Yellow for partners). Joint decision was automatically arbitrated in favor of the decision with higher confidence. Finally, three distinct color-coded feedback messages (participant, partner and joint decision) were displayed.

Participants were instructed to try to maximize the joint accuracy of their social decisions. In order to achieve joint benefit, confidence should be expressed such that the decision with higher probability of correct outcome dominates[6]. For this to happen, the participant needs to factor in the partner’s behavior and adjust her confidence accordingly. For example, if the participant believes that her decision is highly likely to be correct, her confidence should be expressed such that joint decision is dominated by the partner only if the probability that the partner’s decision is correct is even higher (and not, for example, if the partner expressed a high confidence habitually). This social modulation of one’s confidence in a perceptual decision comprises the core of our model of social communication of uncertainty.

Following from an earlier study [19], for each block the participants were led to believe that they were paired with a new, anonymous human partner. In reality, in separate blocks, they were paired with 4 computer generated partners (henceforward, CGPs; see Methods) constructed and tuned to parameters obtained from the participant’s own behavior in the Isolated session: 1: High accuracy and High confidence (HAHC; i.e., this CGP’s decisions were more likely to be more confident as well as more accurate); 2: High accuracy and Low confidence (HALC); 3: Low accuracy and High confidence (LAHC) and 4: Low accuracy and Low confidence (LALC) (see Methods for details). For study 2, we used 2 CGPs (HCA and LCA) while the agent accuracy was similar to those of participants[37] (Wilcoxon ranksum, p=0.37, df=29, zval=0.89). See figure 1-figure supplement 1 for confidence and accuracy data of CGPs. Each participant completed 4 blocks of 200 trials cooperating with a different CGP in each block. Our questionnaire results also confirmed that none of the subject suspected their partners as an artificial one.

Having observed the confidence matching effect in both studies (figure 1.b), a permutation analysis confirmed that this effect did not arise trivially from mere pairing with any random partner[19] (Figure 1-figure supplement 2). The difference between the participant’s confidence and that of their partner was smaller in the Social (vs Isolated) condition (Figure 1-figure supplement 2) consistent with the prediction that participants would match their average confidence to that of their partner in the Social session [19].

Having established the socially emergent phenomenon of confidence matching in the dynamic random dot motion paradigm, we then proceeded to examine choice speed, accuracy and confidence under social conditions (Figure 1.c). We observed that when participants were paired with a high (vs low) confidence partner, there was no significant difference in accuracy between the Social conditions (p=0.92, p=0.75 for study 1 and 2 respectively, Generalized Linear Mixed Model (GLMM), see Supplementary material for details of the analysis (table S1), Figure 1.c top-left panel); confidence, however, was significantly higher (p<0.001 for both studies, table S1, Figure 1.c, middle panel) and reaction times were significantly faster (p<0.001 for both, table S1, Figure 1.c bottom panel) in the HCA vs LCA.

This pattern of dissociations of speed and confidence from accuracy is non-trivial because the expectations of the standard sequential sampling models would be that a change in confidence should be reflected in change in accuracy[22], [23]. Many alternative mechanistic explanations are, in principle, possible. The rich literature on sequential sampling models in the random-dot paradigm permit articulating the components of such intuitive explanations as distinct computational models and comparing them by formal model comparison (see further below).

In order to assess the impact of social context on the participants’ level of subjective uncertainty and rule out two important alternative explanations of confidence matching, we next examined the pupil data. Several studies have recently established a link between human participants’ state of uncertainty and baseline (i.e., non-luminance mediated) variations in pupil size [19], [38]–[42]. If the impact of social context on confidence were truly reflective of a change in the participant’s belief about uncertainty, then we would expect the smaller pupil size when paired with high (HCA) vs low confidence agent (LCA) indicating lower subjective uncertainty. Alternatively, if confidence matching were principally due to pure imitation [43], [44] or due to some form of social obligation in agreeing with others (e.g. normative conformity [45]) without any change in belief, we would expect the pupil size to remain unaffected by pairing condition under social context. We found that during the inter-trial interval (ITI), pupil size was larger in the blocks where participants were paired with low (vs high) confidence agent (figure 2, GLMM analysis, p<0.01 and p<0.001 for study 1 and 2 respectively, see Supplementary material for details of the analysis; table S2). These results may support our hypothesis that social context shapes the participant’s belief about uncertainty. It is important to bear in mind that pupil dilation has been linked to other factors such as mental effort[46], level of surprise [47], and arousal level [48] as well. These caveats notwithstanding, the patterns of pupil dilation within the time period of inter-trial intervals that are demonstrated and replicated here, are consistent with the hypothesis that participants’ subjective belief was shaped by interactions with differently confident partners.

**Figure 2.**
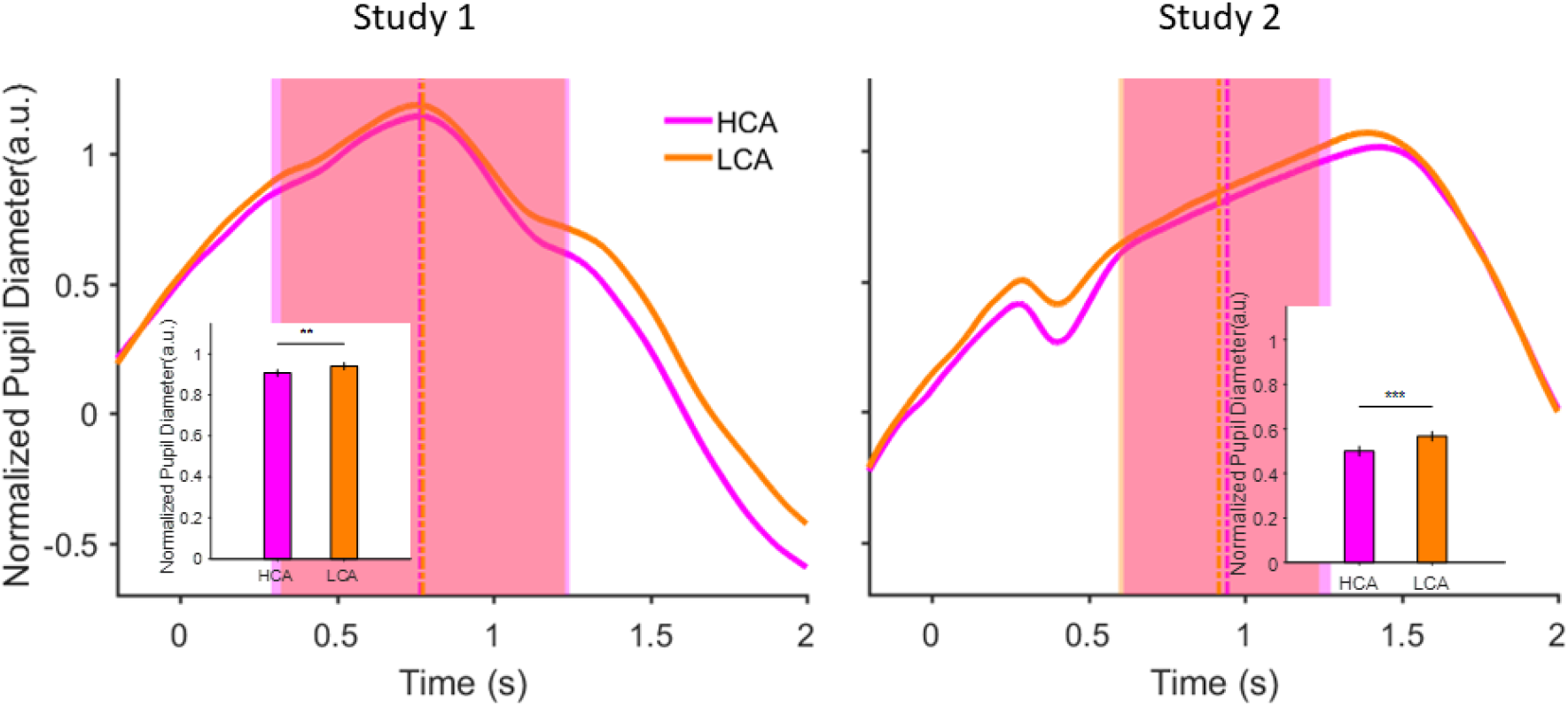
Pupil size during inter trial interval (ITI) under pairing conditions in the social context when participant was paired with a high (HCA) or low confidence (LCA) agent. Normalized pupil diameter aligned to start of ITI period (t=0). Vertical dashed lines show average ITI duration. The shaded areas are one standard deviation of ITI period in each condition. Inset shows grand average (mean) pupil size during ITI under the two Social conditions. Error bars are 95 % confidence interval across trials. (**) indicates p<0.01 and (***) shows p<0.001. In the interest of clarity, signals were smoothed using an averaging filter.

To arbitrate between alternative explanations and develop a neural hypothesis for the impact of social context on decision speed and confidence, we constructed a neural attractor model [49], a variant from the family of sequential sampling models of choice under uncertainty [50]. Briefly, in this model, noisy sensory evidence was sequentially accumulated by two competing mechanisms (red and blue in Figure 3a left) that raced towards a common predefined decision boundary (Figure 3a right) while mutually inhibiting each other. Choice was made as soon as one mechanism hit the boundary. This model has accounted for numerous observations of perceptual and value-based decision making behavior and their underlying neuronal substrates in human[51], and non-human primate [38] brain. Following previous works [38], [52]–[54] we defined model confidence as the time-averaged difference between the activity of the winning and losing accumulators (corresponding to the shaded grey area between the two accumulator traces in Figure 3a right, for the model simulation see figure 3-figure supplement 2) during the period of stimulus presentation (from 0 to 500ms). Importantly, this definition of confidence is consistent with recent findings that computations of confidence continue *after* a decision has been made as long as sensory evidence is available[31], [52], [55], [56]. We also demonstrate that our results do not depend on this specific formulation and also replicate with another alternative method [57](see figure 3-figure supplement 3)

**Figure 3.**
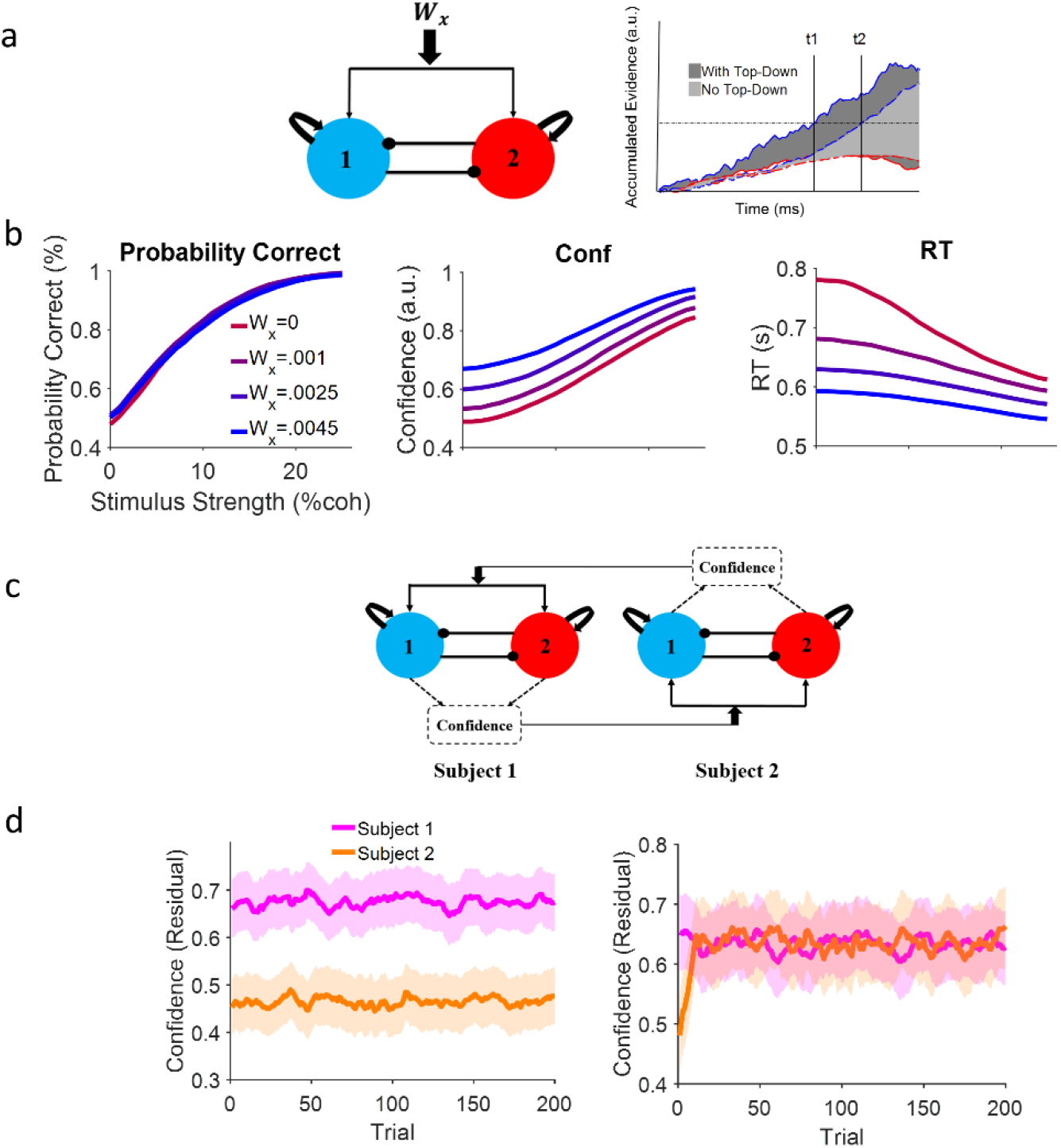
Neural attractor model. (a) Left: A common top-down (*W*_*x*_) current drives both populations, each selective for a different choice alternative. Right: A schematic illustration of the impact of a positive top-down drive on accumulator dynamics. Confidence corresponds to the shaded area between winning (blue) and losing (red) accumulators. Solid lines and dark grey shade: positive top-down drive; dashed lines and light grey shade: zero top-down drive. With positive top-down current, the winner hits the bound earlier (*t1* vs *t2*) and the surface area between the competing accumulator traces is larger (dark vs light grey). (b) Systematic examination of the impact of *W*_*x*_ on model behavior. Left panel: accuracy does not depend on the top down current but confidence (middle) and RT (right) change accordingly. Colors indicate different levels of top-down current. Each curve is the average of 10,000 simulations of the model given the top-down current (c) Dynamic coupling in simulated dyadic interaction. Virtual dyads were constructed by feeding one model’s confidence in previous trial to the other model as top-down drive and vice versa. (d) Left: Unconnected virtual dyad members (*W*_*x*_ = 0) simulate the Isolated condition. As expected, confidence matching is not observed even though the pair receive the exact same sequence of stimuli. Right: When the virtual dyad members are connected with top-down drive proportional to one another’s confidence in previous trial, dyad members’ confidence converge over time. Shadowed areas of the confidence interval 95% resulted from 50 parallel simulations and curves were smoothed by an averaging filter for clearer illustration. The correlation with coherence has been removed from the confidence values via residual analysis (see figure 3-figure supplement 1 confidence values).

Earlier works that demonstrated the relationship between decision uncertainty and pupil-related, global arousal state in the brain[41], [42] guided our modelling hypothesis. Modelling the social context as a global, top-down additive input (Figure 3a; *W*_*x*_) in the attractor model that equally and positively drove both accumulator mechanisms captured our key behavioral observations. The intuitive description of the impact of this global top-down input is illustrated in Figure 3a right: with a positive top-down drive (W_x_>0), the winner (thick blue) and the loser (thick red) traces rise faster and there is a larger (dark grey) surface area between them compared to zero top-down drive (dotted lines and light grey shading); hence, with this positive top-down drive, model decisions (figure 3b) are faster and more confident without any change in accuracy, consistent with our behavioral findings comparing HCA vs LCA conditions (Figure 1c).

We formally compared our model to three alternative, plausible models of how social context may affect the decision process. Without loss of generality, we used data from study 2 to fit the model. The first model hypothesized that partner’s confidence dynamically modulated the decision bound[52] (parameter B in Equation S7). In this model, the partner’s higher confidence reduced the threshold for what counted as adequate evidence, producing the faster RTs under HCA (Figure 1.c). The second model proposed that partner’s confidence changed non-decision time (NDT) [58] (equation S8). Here, pairing with high confidence partner would not have any impact on perceptual processing but instead, non-specifically decrease reaction times across all coherence levels without affecting accuracy. Finally, in the third model, the stimulus-independent perceptual gain[40], [59] parameter of input current (parameter μ_0_ in equation S9) was modulated by partner confidence. Here, higher partner confidence increased the perceptual gain (as if increasing the volume of radio) leading to increased confidence and decreased RT (figure 1.c) and would be consistent with the pupillometry results. In each model, in the Social condition, the parameter of interest was linearly modulated by the confidence of the partner in the previous trial (Importantly, we showed that subjects’ behaviors are dependent to the confidence of the previous trial which validate our trial-by-trial modulation claim, see figure 1-figure supplement 3). Formal model comparison showed that our top-down additive current model was superior to all three alternatives (see figure 3-figure supplement 4).

Having shown that a common top-down drive can qualitatively reproduce the impact of social context on speed-accuracy-confidence and quantitatively excel other alternatives in fitting the observed behavior, we then used the winning model to simulate our interactive social experiment virtually (Figure 3.c). We simulated one decision maker with high confidence (subject 1 in Figure 3.d) and another one with low confidence (subject 2). To simulate subject 1, we slightly increased the excitatory and the inhibitory weights. The opposite was done to simulate subject 2 (see Methods for details). We then paired the two simulated agents by feeding the confidence of each virtual agent (from trial *t-1*)[19] as top-down input to the other virtual agent (in trial *t*).

Using this virtual social experiment, we simulated the dyadic exchanges of confidence in the course of our experiment and drew a prediction that could be directly tested against the empirical behavioral data. Without any fine tuning of parameters or any other intervention, confidence matching emerged spontaneously when two virtual agents with very different confidence levels in Isolated condition (Figure 3.d left) were paired with each other as a dyad (Figure 3.d right).

To identify the neural correlates of interpersonal alignment of belief about uncertainty, we note that previous works using non-invasive electrophysiological recordings in humans engaged in motion discrimination[60], [61] have identified the signature, accumulate-to-bound neural activity characteristic of evidence accumulation in the sequential sampling process. Specifically, these findings show a centropareital positivity (CPP) component in the event-related potential that rises with sensory evidence accumulation across time. The exact correspondence between the neural CPP and elements of the sequential sampling process are not yet clear[61]. For example, CPP could have resulted from the spatial superposition of the electrical activity of both accumulators or be the neural activity corresponding to the difference in accumulated evidence. These caveats notwithstanding, consistent with the previous literature, we found that in the Isolated condition, our data replicated those earlier findings: Figure 4a shows a clear CPP event-related potential whose slope of rise was strongly modulated by motion coherence (GLMM, p<0.001 and p=.01 for study 1 and 2 receptively, see table S5 and figure 4-figure supplement 2 for more details).

**Figure 4.**
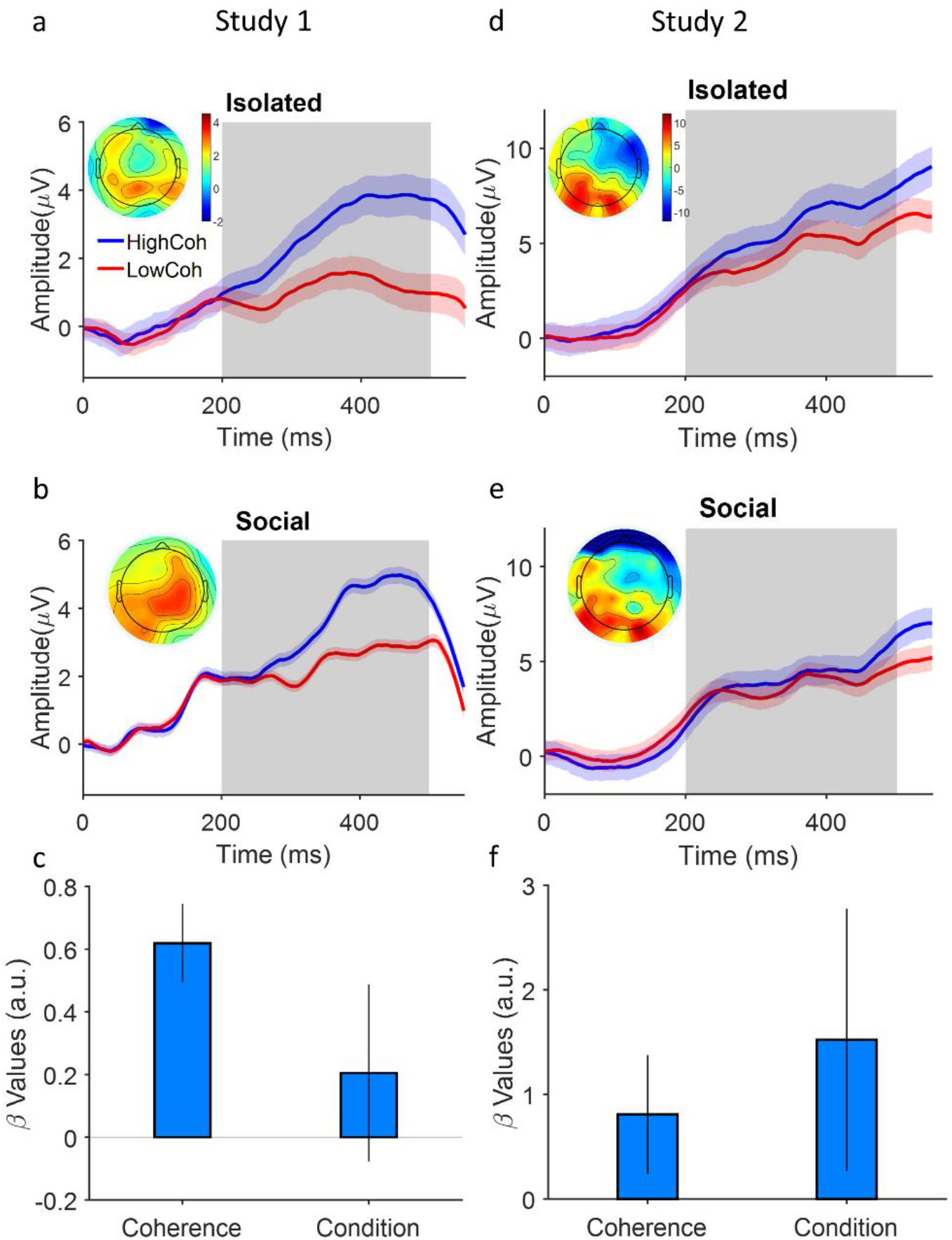
Coupling of neural evidence accumulation to social exchange of information. (a) Centro-Parietal Positivity component (CPP) in the Isolated Condition: event related potentials are time-locked to stimulus onset, binned for high and low levels of coherency (For study 1, Low: 3.2%, 6.4%, 12.8% High: 25.6% and 51.2%. For study 2 (d), Low: 1.6%, 3.2%, 6.4% High: 12.8%, 25.6%) and grand averaged across Centro-patrial electrodes (see methods). Inset shows the topographic distribution of the EEG signal averaged across the time window indicated by the grey area. (b) CPP under Social condition. Conventions the same as panel a. (c) a GLMM model showed the significant relation of Centro-Parietal signals to levels of coherency and Social condition (HCA vs LCA). Error bars are 95% confidence interval over the model’s coefficient estimates. Signals were smoothed by an averaging filter; shaded areas are SEM across trials.

Our model hypothesized that under Social condition, a top-down drive – determined by the partner’s communicated confidence in the previous trial – would modulate the rate of evidence accumulation (cf. Fig 3a). We tested if the CPP slope were larger *within* every given coherence bin when the participant was paired with a high (vs low-) confidence agent. Indeed, the data demonstrated a larger slope of CPP rise under HCA vs LCA (Figure 4c, study 1 for the Social condition p=0.15 but for the second study p<0.01, see table S3 for more details). These findings are the first neurobiological demonstration of interpersonal alignment by coupling of neural evidence accumulation to social exchange of information (for investigation of this coupling in our computational model, see figure 4-figure supplement 3). Importantly, the behavioral confidence matching (Figure 1b) and the pupil data (Figure 2) indicate that this neural-social coupling is associated with the construction of a shared belief about uncertainty.

Second, we note that human medial prefrontal cortex is tonically upregulated during communicative interactions[62] and is a causally necessary neural substrate[63] for constructing a shared cognitive space between interacting agents[8]. Moreover, this brain area has been implicated in perceptual metacognition and neural computations involved in decision confidence[3], [64]. Putting these together with our hypothesis, we predicted (Figure 5a) that a top-down signal from prefrontal cortex (i.e., *W*_*x*_) should drive the centro-parietal (CP) cortex under social condition and, specifically, this top-down drive should be stronger under HCA (vs LCA) condition i.e., when the participant is paired with the high confidence partner. We assessed the effective connectivity (Figure 5b) from prefrontal (PF) to Centro-Parietal (CP) cortex (see Methods) by estimating transfer entropy which is an information theoretic measure quantifying the unidirectional flow of information from PF to CP [65], [66]. We observed stronger flow of information from PF to CP under HCA vs LCA (Figure 5b); importantly, this top-down flow of information was persistent extending from 200ms before the stimulus onset to stimulus presentation [0-500] ms (figure 5-figure supplement 1). This latter observation is consistent with our model’s assumption about constant current (Figure 4a) and in line with a previous report of persistent upregulation of PF activity during social communication[62].

**Figure 5.**
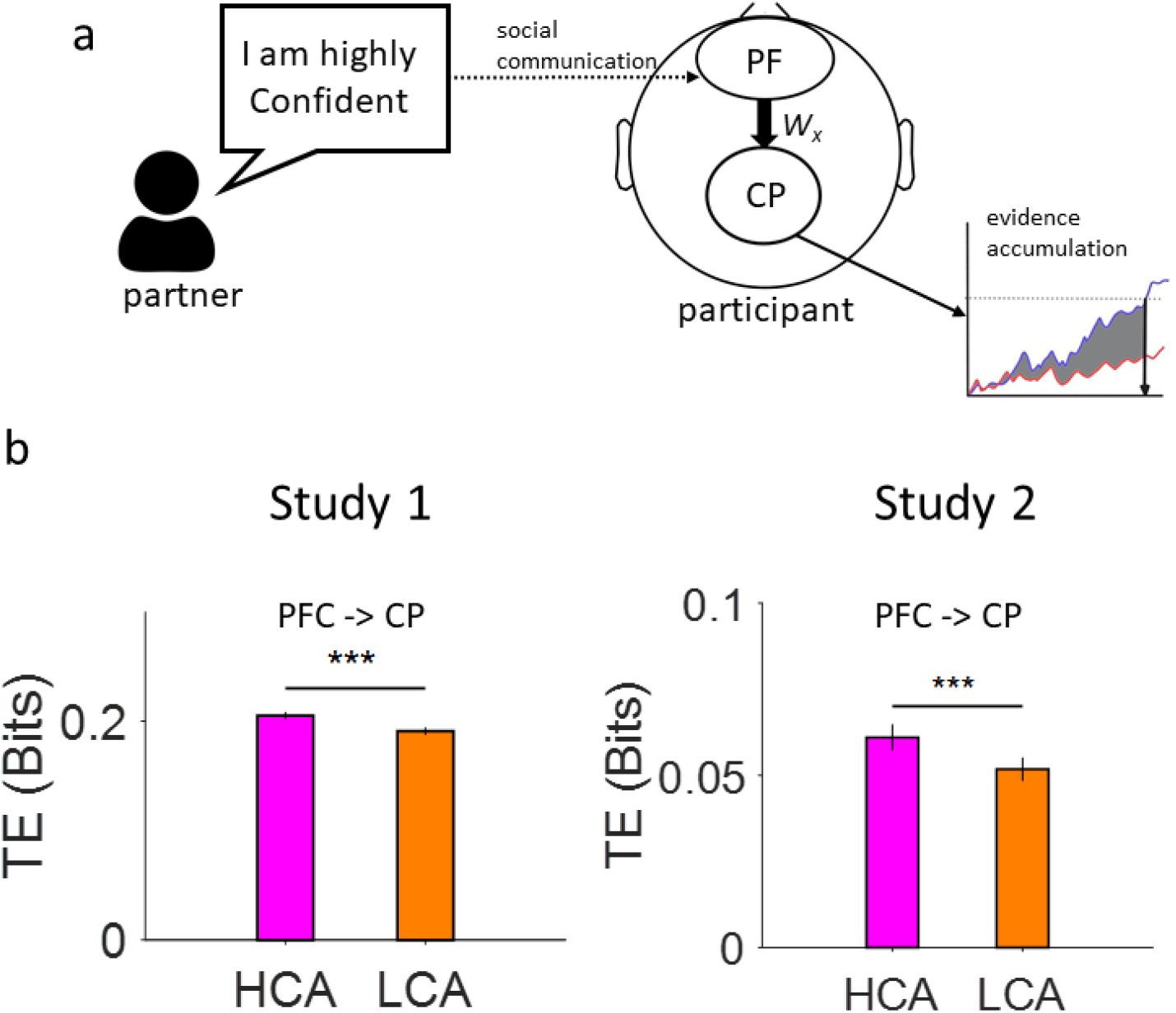
Effective connectivity analysis. (a) A heterogeneous connectivity hypothesis for unifying the social and neurobiological substrates of interpersonal alignment of evidence accumulation in joint decision making under uncertainty. Socially communicated uncertainty (dotted arrow) shapes a top-down drive (*W*_*x*_) from Prefrontal to Centro-Parietal cortex where evidence accumulation takes place. (b) Transfer entropy (TE) estimation under Social (HCA, LCA) conditions. Bar plot shows average of transfer entropy (TE) estimation in the stimulus presentation period of [0-500 ms]. Error bars are Confidence interval 95% across trials. (***) indicates p<0.001; also see Supplementary material table S4 for GLMM).

## Discussion

We brought together two so-far-unrelated research directions: confidence in decision making under uncertainty and interpersonal alignment in communication. Our approach offers solutions to important current problems in each.

For decision science, we provide a model-based, theoretically grounded neural mechanism for going from individual, idiosyncratic representations of uncertainty[2], [3] to socially transmitted confidence expressions[6], [19] that are seamlessly shared and allow for successful cooperation. The social-to-neuronal coupling mechanism that we borrowed from the communication literature[16], [17] is crucial in this new understanding of the neuronal basis of relationship between subjectively private and socially shared uncertainty.

For communication science, by examining perceptual decision making under uncertainty in social context, we created a laboratory model in which the goal of communication was to arrive at a shared belief about uncertainty (rather than creating a look-up table for the meaning of actions[8], [9], [11]). In this way, we could employ the extensive theoretical, behavioural and neurobiological body of knowledge in decision science [20]–[29], [31], [32], [34]–[36], [38], [40], [42] to construct a mechanistic neural hypothesis for interpersonal alignment.

Over the past few years, the efforts to understand the “brain in interaction” have picked up momentum[17], [67]. A consensus emerging from these works is that, at a conceptual level, successful interpersonal alignment entails the mutual construction of a shared cognitive space between brains[8], [17], [68]. This would allow interacting brains to adjust their internal dynamics to converge on shared beliefs and meanings [16], [69]. To identify the neurobiological substrates of such shared cognitive space, brain-to-brain interactions need to be described in terms of information flow, i.e. the impact that interacting partners have on one another’s brain dynamics[17].

The evidence for such information flow has predominantly consisted of demonstrations of alignment of brain-to-brain activity (i.e., synchrony at macroscopic level e.g. fMRI BOLD signal) when people process the same (simple or complex) sensory input [10]–[13], [70] or engage in complimentary communicative [9] roles to achieve a common goal. More recently, dynamic coupling (rather than synchrony) has been suggested as a more general description of the nature of brain-to-brain interaction[16]. Going beyond the intuitive notions of synchrony and coupling, to our knowledge, no computational framework – grounded in the principles of neural computing – has been offered that could propose a plausible quantitative mechanism for these empirical observations of brain-to-brain coupling.

Combining 4 different methodologies, the work presented here undertook this task. Behaviorally, our participants engaged in social perceptual decision making under various levels of sensory and social uncertainty[6], [19]. Emergence of confidence matching (Figure 1.b) showed that participants coordinated their decision confidence with their social partner. Pupil data (Figure 2) suggested that participant’s belief about uncertainty was indeed shaped by the social coordination. A dissociation (Figure 1.c) of decision speed and confidence from accuracy was reported that depended on the social context. This tradeoff, as well as the emergence of confidence matching, were successfully captured by a neural attractor model (Figure 3) in which two competing neural populations of evidence accumulators – each tuned to one choice alternative – were driven by a common top-down drive determined by social information. This model drew predictions for behavior (Figure 3.d) and neuronal activity (Figure 4-5) that were born out by the data. Social exchange of information modulated (1) the neural signature of evidence accumulation in the parietal cortex and (2) a sustained top-down flow of information from prefrontal to parietal cortex.

Although numerous previous works have employed sequential sampling models to explain choice confidence, the overwhelming majority [22], [23], [25]–[32] have opted for the drift diffusion family of models. Neural attractor models have so far been rarely used to understand confidence[53], [54], [71]. Our attractor model is a reduced version[49] of the original biophysical neural circuit model for motion discrimination[71]. The specific affordances of attractor models allowed us to implement social context as a sustained, tonic top-down feedback to both accumulator mechanisms. More importantly, we were able to simulate social interactive decision making by virtually pairing any given two instances of the model (one for each member of a dyad) with each other: the confidence produced by each in a given trial served as top-down drive for the other in the next trial. Remarkably, a shared cognitive space about uncertainty (i.e., confidence matching) emerged spontaneously from this simulated pairing without us having to tweak any model parameters.

At a conceptual level, deconstructing the social communication of confidence into a comprehension and a production process [9] is helpful. Comprehension process refers to how socially communicated confidence is incorporated in the recipient brain and affects their decision-making. Production process refers to how the recipient’s own decision confidence is constructed to be, in turn, socially expressed. It is tempting to attribute the CPP neural activity in the parietal cortex to the production process. Comprehension process, in turn, could be the top-down feedback from prefrontal brain areas previously implicated in confidence and metacognition [3], [21], [72] to the parietal cortex. However, we believe that our neural attractor model in particular and the empirical findings do not lend themselves easily to this conceptual simplification. For example, the evidence accumulation process can be a part of the production (because confidence emerges from the integrated difference between accumulators) as well as the comprehension process (because the rate of accumulation is modulated by the received social information). As useful as it is, the comprehension/production dichotomy’s limited scope should be recognized. Instead, armed with the quantitative framework of neural attractor models (for each individual) and interactive virtual pairing (to simulate dyads), future studies can now go beyond the comprehension/production dichotomy and examine the neuronal basis of interpersonal alignment with a model that have a strong footing in biophysical realities of neural computation.

Several limitations apply to our study. We chose different sets of coherence levels for the discovery (experiment 1) and replication (experiment 2). This choice was made deliberately. In experiment 1 we included a very high coherence (51%) level to optimize the experimental design for demonstrating the CPP component in the EEG signal. In experiment 2, we employed peri-threshold coherence levels in order to focus on behavior around the perceptual threshold to strengthen the model fitting and model comparison. This trade-off created some marginal differences in the observed effect sizes in the neural data across the two studies. The general findings were in good agreement.

Moreover, due to the inherent limitations of EEG methodology and its poor spatial resolution, it was not possible to limit the connectivity analysis to (or draw a precise conclusion about) the exact anatomical location of our hypothesized neural origin of the top-down signal in the prefrontal cortex. Moreover, our study did not directly examine neural alignment between interaction partners. We measured the EEG signal one participant at a time. The participant interacted with an alleged (experimenter-controlled) partner in any given trial. Our experimental design, however, permitted strict experimental control and allowed us to examine the participants’ social behavior (i.e., choices and confidence), pupil response and brain dynamics as they achieved interpersonal alignment with the partner. Moreover, while the hypotheses raised by our neural attractor model did examine the nature of brain dynamics involved in evidence accumulation under social context, testing these hypotheses did not require hyper-scanning of two participants at the same time. We look forward to future studies that use the behavioral and computational paradigm described here to examine brain-to-brain neural alignment using hyper-scanning.

## Materials and Methods

### Participants

A total of 27 participants (12 in Experiment 1 and 15 in Experiment 2; 10 females; average age: 24 years; all naïve to the purpose of the experiment) were recruited for a two-session experiment – Isolated and Social session. All subjects reported normal or corrected-to-normal vision. The participants did several training sessions in order to become familiar with the procedure and reach a consistent pre-defined level of sensitivity (see Methods for more details).

**Recruitment**

Participants volunteered to take part in the experiment in return for course credit for study 1. For study 2, a payment of 80,000 Toman equivalent to 2.5€ per session was made to each participant. On the experiment day, participants were first given the task instructions. Written informed consent was then obtained. The experiments were approved by the local Ethics Committee at Shaheed Rajaei University’s Department of computer engineering.

### Task Design

In the Isolated session, each trial started with a red fixation point in the center of the screen (diameter 0.3°). Having fixated for 300ms (in study 1, for a few subjects with eye monitoring difficulty this period shortened), two choice-target points appeared at 10° eccentricity corresponding to the two possible motion directions (left and right) (figure 1). After a short random delay (200-500 ms, truncated exponential distribution), a dynamic random dot motion stimulus was displayed for 500ms in a virtual aperture (5° diameter) centered on the initial fixation point. These motion stimuli have been described in detail elsewhere[35]. At the end of the motion stimulus a response panel (see Figure 1a) was displayed on the screen. This response panel consisted of a horizontal line extending from left to the right end of the display, centered on the fixation cross. On each side of the horizontal line, 6 vertical rectangles were displayed side by side (Figure 1a) corresponding to 6 confidence levels for each decision alternative. The participants reported the direction of the random dot motion stimulus and simultaneously expressed their decision and confidence using the mouse.

The rectangles on the right and left of the midpoint corresponded to the right and left choices, respectively. By clicking on the rectangles further the midpoint participants indicated higher confidence. In this way, participant indicated their confidence and choice simultaneously[28], [73] For Experiment 1, response time was defined as the moment that the marker deviated (more than one pixel) from the center of the screen. However, in order to rule out the effect of unintentional movements, for the second study we increased this threshold to one degree of visual angle. The participants were informed about their accuracy by a visual feedback presented in the center of the screen for 1 second (Correct or Wrong).

In the Social session, the participants were told they were paired with an anonymous partner. In fact, they were paired with a Computer-Generated Partner (CGP) tailored to the participant’s own behavior in their Isolated session. The participants did not know about this arrangement. Stimulus presentation and private response phase were identical to the Isolated session. After the private response, the participants were presented with a social panel right (figure 1). In this panel, the participant’s own response (choice and confidence) were presented together with that of their partner for 1 second. The participant and the partner responses were color-coded (White for participants; Yellow for partners). Joint decision was determined by the choice of the more confident person and displayed in green. Then, three distinct color-coded feedbacks were provided.

In both Isolated and Social sessions, the participants were seated in an adjustable chair in a semi-dark room with chin and forehead supported in front of a CRT display monitor (First study: 17 inches; PF790; refresh rate, 85 Hz; screen 164 resolution, 1024 × 768; viewing distance, 57 cm, second study: 21 inches; Asus VG248; refresh rate, 75 Hz; screen resolution, 1024 × 768; viewing distance, 60 cm). All the code was written in PsychToolbox [74]–[76].

**Training procedure**

Each participant went through several training sessions (on average 4) to be trained on Random Dot Motion (RDM) task. They first trained in a response free (i.e., Reaction Time) version of the RDM task in which motion stimulus was discontinued as soon as the participant responded. They were told to decide about the motion direction of dots as fast and accurately as possible[28]. Once they reached a *stable* kevel of accuracy and RT, they proceeded to the main experiment. Before participating in the main experiment, they performed another 20-50 trials of warm-up. Here the stimulus duration was fixed and responses included confidence report. For the social sessions, participants were told that in every block of 200 trials, they would be paired with a different person, seated in another room, with whom they would collaborate. They were also instructed about the joint decision scheme and were reminded that the objective in the social task was to maximize collective accuracy. Data from training and warm-up trials were included in the main analysis.

### Procedure

Each participant performed both the Isolated and the Social task. In the Isolated session, they did one block containing 200 trials. Acquired data were employed to construct four computer partners for the first study and two partners for the second study. We used the procedure introduced in a previous works to generate CGPs[19], [77]. In the first study, the four partners were distinguished by their level of average accuracy and overall confidence: High accuracy and High confidence (HAHC), High accuracy and Low confidence (HALC), Low accuracy and High confidence (LAHC) and finally Low accuracy and Low confidence (LALC). For the second study partners only differed in confidence: High confidence agent (HCA) and low confidence agent (LCA). Each participant performed one block of 200 trials for each of the paired partners – 800 overall for study 1 and 400 overall for study 2.

In the Social session, participants were told to try to maximize the joint decision success [19]. They were told that their payment bonus depended on by their joint accuracy[37]. While performing the behavioral task, EEG signals and pupil data were also recorded.

### Computer Generated Partner

In study 1, following [19], four partners were generated for each participant tuned to the participant’s own behavioral data in the Isolated session. Briefly, we created 4 simulated partners by varying their mean accuracy (high or low) and mean confidence (high or low). First, in the isolated session, the participant’s sensory noise (σ) and a set of thresholds that determined the distribution of their confidence responses were calculated (see Methods also). Simulated partner’s accuracy was either high (0.3xσ) or low (1.2xσ). Mean confidence of simulated partners were also set according to the participant’s own data. For low confidence simulated partner, average confidence was set to the average of participant’s confidence in the low coherence (3.2% and 6.4%) trials. For the high confidence simulated partners, mean confidence was set to the average confidence of the participant in the high coherence (25.6% and 51.2%) trials. Reaction times were chosen randomly by sampling from a uniform random distribution (from 0.5 to 2 seconds). Thus, in some trials the participant needed to wait for the partner’s response.

Having thus determined the parameters of the simulated partners, we then generated the sequence of trial-by-trial responses of a given partner using the procedure introduced by Bang et al. [19]. To produce the trial by-trial responses of a given partner, we first generated a sequence of coherence levels with given directions (+ for rightward and – for leftward directions). Then we created a sequence of random values (sensory evidence), drawn from a Gaussian distribution with mean of coherence levels and variance of σ (sensory noise). Then, via applying the set of thresholds taken from the participant’s data in Isolated condition, we mapped the sequence of random values into trial-by-trial responses to generate a partner with a given confidence mean. Finally, to simulate lapses of attention and response errors, we randomly selected a response (from a uniform distribution over 1–6) on 5% of the trials (See figure 1-figure supplement 1 for the accuracy and confidence of the generated partners)

For study 2, we used the same procedure as study 1 and simulated two partners. These partners accuracy was similar to the participant but each had a different confidence means (High confidence and low confidence partners). Therefore, we kept the *σ* constant and only change the confidence. For low confidence simulated partner, average confidence was set to the average of participant’s confidence in the low coherence (1,6%, 3.2% and 6.4%) trials. For the high confidence simulated partners, mean confidence was set to the average confidence of the participant in the high coherence (12.8% and 25.6%) trials.

**Signal detection theory model for isolated sessions**

In study 1 and 2, we simulated 4 and 2 artificial partners, respectively. We followed the procedure described by Bang et al [19]. Briefly, working with the data from the isolated session, the sensory noise (*σ*) and response thresholds (***θ***) for each participant were calculated using a signal detection theory model. In this model, the level of sensory noise (*σ*) determines the participant’s sensitivity and a set of 11 thresholds determines the participant’s response distribution, which indicate both decision (via its sign) and confidence within the same distribution (see below).

On each trial, the sensory evidence, *x*, is sampled from a Gaussian distribution, *x* ∈*N*(*s*, σ^2^). The mean, *s*, is the motion coherence level and is drawn uniformly from the set *s* ∈ *S* = {−.512, −.256, −.128, −.064, −.032, .032, .064, .128, .256, .512} (for the second study *S* = {−.256, −.128, −.064, −.032, -.016, .016 .032, .064, .128, .256}). The sign of *s* indicates the correct direction of motion (right = positive) and its absolute value indicates the motion coherency. The standard deviation, σ, describes the level of sensory noise and is the same for all stimuli. We assumed that the internal estimate of sensory evidence (z) is equal to the raw sensory evidence (x). If z is largely positive, it denotes high probability of choosing right direction and vice versa for largely negative values.

To determine the participant’s sensitivity and the response thresholds, first, we calculated the distribution of responses (*r*, ranging from -6 to 6, where the participant’s confidence was (*c* = |*r*|), and her decision was determined by the sign of *r*). Equation S1 shows the response distribution.

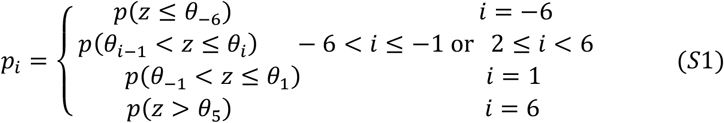

Using ***θ*** and *σ*, we mapped z to participants response (*r*). We found thresholds *θ*_i_ over S where *i* = −6, −5, -4, -3, -2, −1,1,2, 3, 4,5 such that:

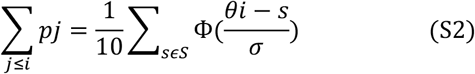

where Φ is the Gaussian cumulative density function. For each stimulus, *s*∈*S*, the predicted response distribution, *p*(*r* = *i*|*s*) calculated by S3:

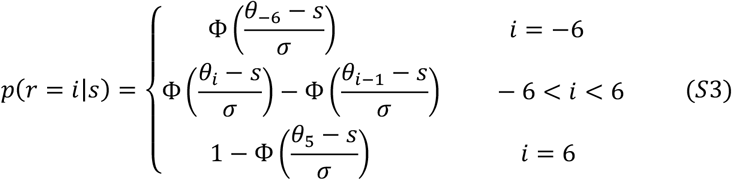

From here, the model’s accuracy could be calculated by S4:

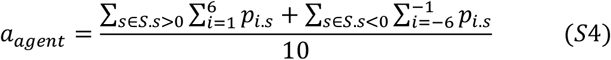

Given participant’s accuracy, we could find a set of ***θ*** and *σ*.

**Confidence estimation**

Once we had determined ***θ*** and *σ*, we could produce a confidence landscape with a specific mean. In order to generate one high confidence and another low confidence partner, we needed to alter mean confidence by modifying the ***θ***. There could be an infinite number of confidence distribution with the desired mean. We were interested in the maximum entropy distribution that satisfied two constraints: mean confidence should be specified, and the distribution must sum to

1. Using Lagrange multiplier (*λ*) the response distribution was calculated as:

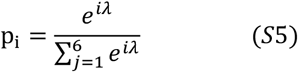

with *λ* chosen by solving the constraint

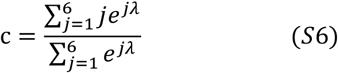

We transformed confidence distributions (1 to 6) to response distributions (−6 to -1 and 1 to 6) by assuming symmetry around 0. Figure S1 shows the accuracy and confidence of generated agents.

### Computational Model

We employed a previously described attractor network model[49] which is itself the reduced version of an earlier one[71] inspired by the mean field theory. The model consists of two units simulating the average firing rates of two neural populations involved in information accumulation during perceptual decisions (figure 3a). When the network is given inputs proportional to stimulus coherence levels, a competition breaks out between two alternative units. This race would continue until firing rates of one of the two units reaches the high-firing-rate attractor state at which point the alternative favored by the unit is chosen. The details of this model have been comprehensively described elsewhere[49].

Each unit was selective to one choice (1,2) and received an input as follow:

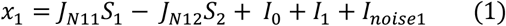

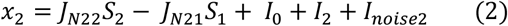

Where *J*_*N11*_ and *J*_*N22*_ indicated the excitatory recurrent connection of each population and *J*_*N12*_ and *J*_*N21*_ showed the mutual inhibitory connection values. For the simulation in figure 3.b we set the recurrent connections to 0.3157 nA and inhibitory ones to 0.0646 nA. *I*_*0*_ indicated the effective external input which was set to 32.55 nA. *I*_*noise1*_/*I*_*noise2*_ stood for the internal noise in each population unit. This zero mean Gaussian white noise was generated based on the time constant of 2 ms and standard deviation of 0.02 nA. *I*_*1*_/*I*_*2*_ indicated the input currents proportional to the motion coherence level such that:

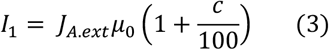

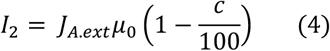

Where *J*_*A*.*ext*_ was the average synaptic coupling from the external source and set to 0.0002243 (nA.Hz^-1^), *c* was coherence level and *µ*_0_, a.k.a perceptual gain, was the input value when the coherence was zero (set to 45.8 Hz).

*S*_*1*_ and *S*_*2*_ were variables representing the synaptic current of either population and were proportional to the number of active NMDA receptors. Whenever the main text refers to accumulated evidence, we refer to *S*_*1*_ and *S*_*2*_ variables. Dynamics of these variables were as follow:

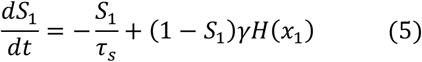

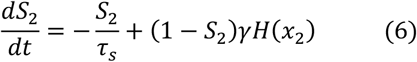

Where τ*s*, the NMDA receptor delay time constant, was set to 100 ms, γ set to 0.641 and the time step, *dt*, was set to 0.5 ms. Dynamical equations 5 and 6 were solved using forward Euler method[49]. (H), the generated firing rates of either populations, was calculated by:

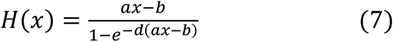

Where *a, b*, and *d* were set to 270 Hz nA^-1^, 108 Hz and 0.154 s, respectively. These constants indicated the input-output relationship of a neural population.

The model’s choice in each trial was defined as the accumulated evidence of either population that first touched a threshold, and the decision time was defined as the time when the threshold was touched. Notably, the decision threshold was set to *S*_*threshold*_ = 0.32. Moreover, the confidence was defined as the area between two accumulators (*S*_*1*_ and *S*_*2*_ in equations 5 and 6), in the time span of 0 to 500 ms, which was defined as:

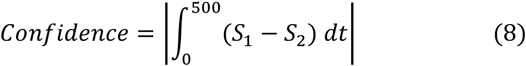

which was normalized by following logistic function[38]:

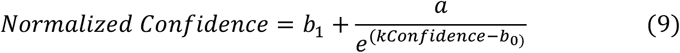

where the values of *b*_1_, *a, k* and *b*_0_ were set to 1.32, -0.99, 5.9 and 0.16 respectively for model on entire trials of subjects in Isolated sessions; *Confidence* is calculated in equation (8) in time period of [0-500] ms.

In line with previous studies, we calculated the absolute difference between accumulators (equation 8) [38], [53]. In this formulation, confidence is calculated from model activity during the stimulus duration[54]. Notably, in our confidence definition, we integrated the accumulators’ difference even when the winning accumulator hit the threshold (post-decision period) [52], [78], [79]. This formulation of confidence provided a successful fit to subjects’ behaviors (figure 3-figure supplement 5). To demonstrate that our key findings do not depend on this specific formulation, we implemented another alternative method [57] and showed qualitatively similar results (figure 3-figure supplement 3) are obtained.

We calibrated the model to the data from the Isolated condition to identify the best fitting parameters that would describe the participants’ behavior in isolation. In this procedure decision threshold, inhibitory and excitatory connections, non-decision time (set 0.27s) and *µ*_0_ were considered as the model variables.

In order to explain the role of social context on participant’s behavior, we added a new input current to the model. Importantly we kept all other parameters of the model identical to the best fit to the participants’ behavior in the Isolated situation:

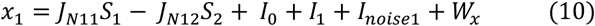

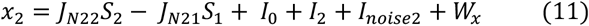

In order to evaluate the effect of *W*_*x*_ on the RT, accuracy and confidence, we simulated the model while systematically varying the values of *W*_*x*_ (figure 3.b).

Having established the qualitative relevance of W_x_ in providing a computational hypothesis for the impact of social context, then we defined W_x_ proportional to the confidence of partner as follows:

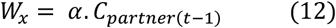

where *t* was the trial number. The model inputs were identical to Isolated situation expect for the top-down current of *W*_*x*_ which indicates the social input where *α* was a normalization factor (or coupling coefficient) and *C*_*partner*(*t*−1)_ indicates the partner’s confidence in the previous trial. Thus, we added a social input based on the linear combination of the partner’s confidence in the previous trial. Notably, the behavioral effect reported in the main script is also evident respect to the confidence of the agent in the previous trial (figure 1-figure supplement 3 and table S6).

For simulations reported in figure 3.d, we created high and low confident models by altering the inhibitory and excitatory connections of the original model. For the high confident model, excitatory and inhibitory connections were set to 0.3392 and 0.0699. For the low confident model excitatory and inhibitory connections were set to 0.3163 and 0.0652 respectively. For the simulation of social interaction (figure 4.f), we coupled two instances of the model using equation (12) with *α* set to - 0.0005 and 0.004 for high confident and low confident models, respectively. We ran the parallel simulations 50 times and reported the average results.

In order to remove the effect of coherence levels from models’ confidence, we measured the residuals of models’ confidence after regressing out the impact of coherence. Using this simple regression model:

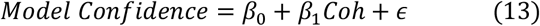

where *Coh* is the motion coherence level and *ϵ* is the error term, we removed the information explainable by motion coherence levels from confidence data as following. Confidence residuals were therefore:

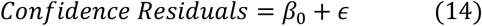

All the simulations of model in the text – and parameters reported in the method-are related to the model calibrated on the collapsed data of all subjects (n=3000 for Isolated sessions of study 2).

**Alternative Formulations for Confidence in the computational model**

In our main model, confidence is formalized by equation 8 in the main text. We calculated the integral of difference between the losing and the winning accumulator during the stimulus presentation. This value would then be fed into a logistic function (equation 9) to produce the final confidence reported by the model (figure 3.b middle panel). To demonstrate the generality of our findings, we used another alternative (but similar) formulation in the previous literature for confidence representation. In Figure 3-figure supplement 3, we compare the resulting “raw” confidence values (i.e., confidence values before they are fed to equation 9).

Alternative formulations for confidence are

1. For comparison we plot our main formulation (equation 8 of main text) in Figure 3-figure supplement 3.a
2. By calculating the difference between winning and losing accumulator at the END of stimulus duration [78] (figure 3-figure supplement 3.b, we call this End method)

Our simulations showed that our formulation (figure 3-figure supplement 3.a) shows an expected modulation to top-down currents. Figure 3-figure supplement 3.b also shows a similar pattern which indicates our results is not different from End method. Therefore, our computational results could be generalized to different confidence representation methods

**Model Comparison**

For model comparison, we used the fitted parameters from the isolated session (Study 2 only without loss of generality). The model parameters for the isolated condition were extracted for each participants in their own respective isolated session (n = 3000 across all participants). Then we compared all “alternative” models with a “single free parameter” to determine the model with the best account to behavioral data in social sessions (n=6000 across all participants). We considered three alternative models for the comparison. Note that in all models *a* is the normalization factor and the free parameter.

Bound Model

We hypothesized that partner’s confidence modulates the participant’s decision boundary according to:

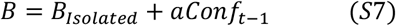

*B* determines the threshold applied on the solution of the equations 5 and 6 (see Methods). *B*_*Isolated*_ denotes the threshold in the isolated model. In this model, in social condition the bound depends on the value of the agent’s confidence in the previous trial. Note that the optimum value of *a*, normalization or coupling factor, is most likely to be negative since it generates lower RTs in social vs isolated situation.

Non-Decision Time (NDT) model

We hypothesized that NDT would be modulated by confidence of agent in the previous trial. Here,

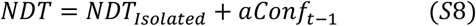

*NDT*_*Isolated*_ was the NDT fitted on the isolated data. Similarly, the optimum *a*, was expected to be negative.

Gain Model

We hypothesized that social information modulated the perceptual gain defined as:

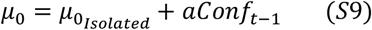

Where *μ*_*0*_ denotes the input value of the model when motion coherence is zero (equation 3 and 4, Methods) and 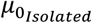 was calculated based on isolated data. If *a* is positive, then *μ*_*0*_ would be greater under social condition vs isolated condition, which in turn generates lower RTs and higher confidence.

In order to incorporate the accuracy, RT and confidence in model comparison, we calculated the RT distribution of trials in each of the 12 confidence levels, six for left decision (−6 to -1) and six for right decision (1 to 6). The RT in each level was further divided into two categories[80] (less than 700 ms and larger than 700 ms). We tried to maximized the likelihood of behavioral RT distribution in each response level (confidence and choice) given the model structure and parameters. The probability matrix was defined as follow:

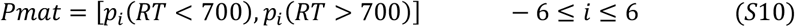

Where *i* is confidence levels ranging from -6 to 6. Note, the probability was calculated based on all trials in our behavioral data set (6000 trials). The model’s probability matrix was also calculated in a similar manner. Hence, we derived a probability matrix of 12 response levels and 2 RT bins. The likelihood function was defined as follows:

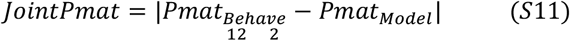

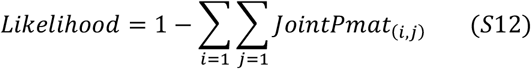

Since we used similar parameters for the models (all models had 1 free parameter, *a*) we could directly compare likelihood values corresponding to each model (figure S10). The model with the highest likelihood is the preferred model; the parameters were found via MATLAB *fminsearch* function.

Additionally, considering the best likelihood for the models confirms the same pattern, lending further support to the superiority of top-down model. Overall, the “Top-Down” model outperformed the three alternative models in explaining behavioral data. The fitted coupling parameter for the top-down model obtained here was 0.0017.

Intuitively, it is possible to see how each model fails to capture the entirety of our empirical results. NDT was able to explain unchanged accuracy and lower RT, but it failed to explain the confidence difference between isolated and social. The bound model also accounted for lower RT in social condition but was unable to explain the pattern of accuracy and confidence. Finally, the gain model could show an increase in confidence and decrease in RT, but stronger gain under social condition, would result in higher accuracy which was different from the behavioral data.

### Eye Monitoring and Pupilometery

In both studies, the Eye movements were recorded by an EyeLink 1000 (SR-Research) device with a sampling rate of 1000Hz which was controlled by a dedicated host PC. The device was set in a desktop and pupil-corneal reflection mode while data from the left eye was recorded. At the beginning of each block, for most subjects, the system was recalibrated and then validated by 9-point schema presented on the screen. One subject was showed a 3-point schema due to the repetitive calibration difficulty. Having reached a detection error of less than 0.5°, the participants were led to the main task. Acquired eye data for pupil size were used for further analysis. Data of one subject in the first study was removed from further analysis due to storage failure.

Pupil data were divided into separate epochs and data from Inter-Trials Interval (ITI) were selected for analysis. Then, blinks and jitters were detected and removed using linear interpolation. Values of pupil size before and after the blink were used for this interpolation. Data was also mid-pass filtered using Butterworth filter (second order, [0.01, 6] Hz)[55]. The pupil data was baseline corrected by removing the average of signal in the period of [-1000 0] ms interval (before ITI onset) and then z-scored. Importantly, trials with ITI>3s were excluded from analysis (365 out of 8800 for study 1 and 128 out 6000 for study 2. also see table S7 and Selection criteria for data analysis in Supplementary Materials)

### EEG Signal recording and preprocessing

For the first study, a 32-channel eWave32 amplifier was used for recording which followed the 10– 10 convention of electrode placement on the scalp (for the locations of the electrodes, see the figure 4-figure supplement 1; right mastoid as the reference). The amplifier, produced by ScienceBeam (http://www.sciencebeam.com/) provided a 1K sampling rate [81]. For the second study we used a 64-channel amplifier produced by LIV team (http://lliivv.com/en/) with 250 Hz sampling rate (see the electrode placement in figure 4-figure supplement 1).

Raw data were analyzed using EEGLAB software[82]. First, data were notch filtered in the range of 45-55 Hz in order to remove the line noise. Using an FIR filter in the range of 0.1-100 Hz, high frequency noise was also removed from data. Artifacts were removed by visual inspection using information from independent component analysis. Noisy trials were removed by also visual inspection. Noisy channels were interpolated using EEEGLAB Software. The signals were divided into distinct epochs aligned to stimulus presentation ranging from 100 ms pre-stimulus onset until 500 ms post-stimulus offset. After preprocessing, EEG data in the designated epochs that had higher (lower) values than 200 (−200) μV were excluded from analysis (see table S7 and Methods for details data analysis) [34]. In the first study, EEG recording was not possible in two participants due to unresolvable impedance calibration problems in multiple channels.

**Relation of CPP to coherence and social condition**

Activities of centroparietal area of the brain is shown to be modulated with coherence level. Here, we showed that CPP activities are statistically related to the coherence levels (Figure 4-figure supplement 2. Top-row) in both studies. Furthermore, we tested how much this relationship is dependent to social condition (HCA, LCA, Figure 4-figure supplement 2. Bottom-row). Our analysis showed that the slope (respect to coherence levels) is different in HCA vs LCA (also see Table S3). Notably, this effect is in line with our neural model prediction (See figure 4-figure supplement 3, next section).

### EEG Analysis of effective connectivity

In order to compare the flow of information flow from Prefrontal to Centro-Partial cortex under the two conditions of social interaction, we used the transfer entropy (TE) method which has been used in effective connectivity analysis[83]. Using the Markov process, Schreiber proposed a measure of information flow between two time series with a specified lag[65]. This measure is called transfer entropy which under specific conditions could be considered as the conditional mutual information [84]. To quantify TE, we used the TIM toolbox (http://www.cs.tut.fi/~timhome/tim/tim.htm). Various lags between the two signals recorded from the Prefrontal and Centro-Partial group of electrodes were used. These lags varied from 10 to 200 ms in order to cover the time span observed for information flow between pareital and frontal cortex[85], [86]. Then, TE measure plotted in Figure 7.b was the grand average of TE [66] over all lags [65]. For this analysis, data were initially filtered using a surface Laplacian (i.e., a spatial transformation also known as current source density) filter[87]. This step is mainly useful as a preparation for connectivity analysis and improves the topological localization and more importantly reduces the impact of volume conduction. We used the exact same procedure and parameters in both studies. Because of difference in the recording devices and sampling frequencies, the results of TE have different scales between study 1 and 2.

**Selection criteria for data analysis**

The data included in both studies could be classified into three main categories: Behavioural, Eye tracking and EEG. For the behavioural analysis, data from all participants were included. In study 1, Eye tracking data from one participant was lost due to storage failure. For pupil analysis, we excluded the trials with inter trial intervals (ITI) longer than 3 seconds (∼ 4% of trials in study 1 and ∼ 2% for study 2).

We also analysed brain data of participants in both studies. For the ERP analysis, we excluded trials with an absolute amplitude greater than 200 microvolts (overall less than 1% for both trials) as this data was deemed as outlier. Moreover, noisy trials and ICA components (around 5% of components in study 2) were rejected by visual inspection. Noisy electrodes were also interpolated (∼ 8% of electrodes in study 2); see table S7 for more details. In study 1, EEG data from 2 participants were lost due to a technical failure. All data (behaviour, eye tracking and EEG) for study 2 were properly stored, saved and made available at https://github.com/JimmyEsmaily/ConfMatch

### Statistical Analysis

For hypothesis testing, we employed a number of generalized linear mixed models (GLMM). Unless otherwise stated, in our mixed models, participant was considered as random intercept. Details of each model is described in tables S1-S6 in the supplementary materials. This approach enabled us to separate the effects of coherency and partner confidence. For RT and confidence, we assumed that the data is normality distributed. For the accuracy data we assumed the distribution is Poisson. We used a maximum likelihood method for fitting. All p-values reported in the text were drawn from the GLMM method, unless stated otherwise.

**Permutation test to confirm confidence matching**

A key null hypothesis (p(ϑ) where ϑ is the measure of interest: confidence matching) that we ruled out was that confidence matching was forced by the experimental design limitations and, therefore, would be observed in any random pairing of participants within our joint decision making setup. To reject this hypothesis, we performed a permutation test following Bang et al [19] (see their Supplementary Figure 3 for further details). For each participant and corresponding CGP pair, we defined |*c*_*1*_-c_2_| where *c*_*i*_ is the average confidence of participant *i* in a given pair. We then estimated the null distribution for this variable by randomly re-pairing the participant with other participants and computing the mean confidence matching for each such re-paired set (total number of sets 1,000). In Figure 1-figure supplement 2 (bottom row), the red line shows the empirically observed mean of confidence matching in our data. The null distribution is shown in black. Proportion of values from the null distribution that were less than the empirical mean was p∼0).

In addition, we defined an index for measuring the confidence matching (figure 1-figure supplement 2. First row): *Δm* = |*C*_*isolated*(*Subject*)_ − *C*_*agent*_ | − |*C*_*social*(*Subject*)_ − *C*_*agent*_|. The larger the *Δm* the higher is the confidence matching. Although we did not observe a significant effect of *Δm*, we showed that this index is significantly different from zero in the HCA condition.

## Acknowledgments

JE and BB were supported by the European Research Council (ERC) under the European Union’s Horizon 2020 research and innovation programme (819040 - acronym: rid-O). BB was supported by the NOMIS foundation and Templeton Religion Trust.

---

**Figure 1-figure supplement 1.**
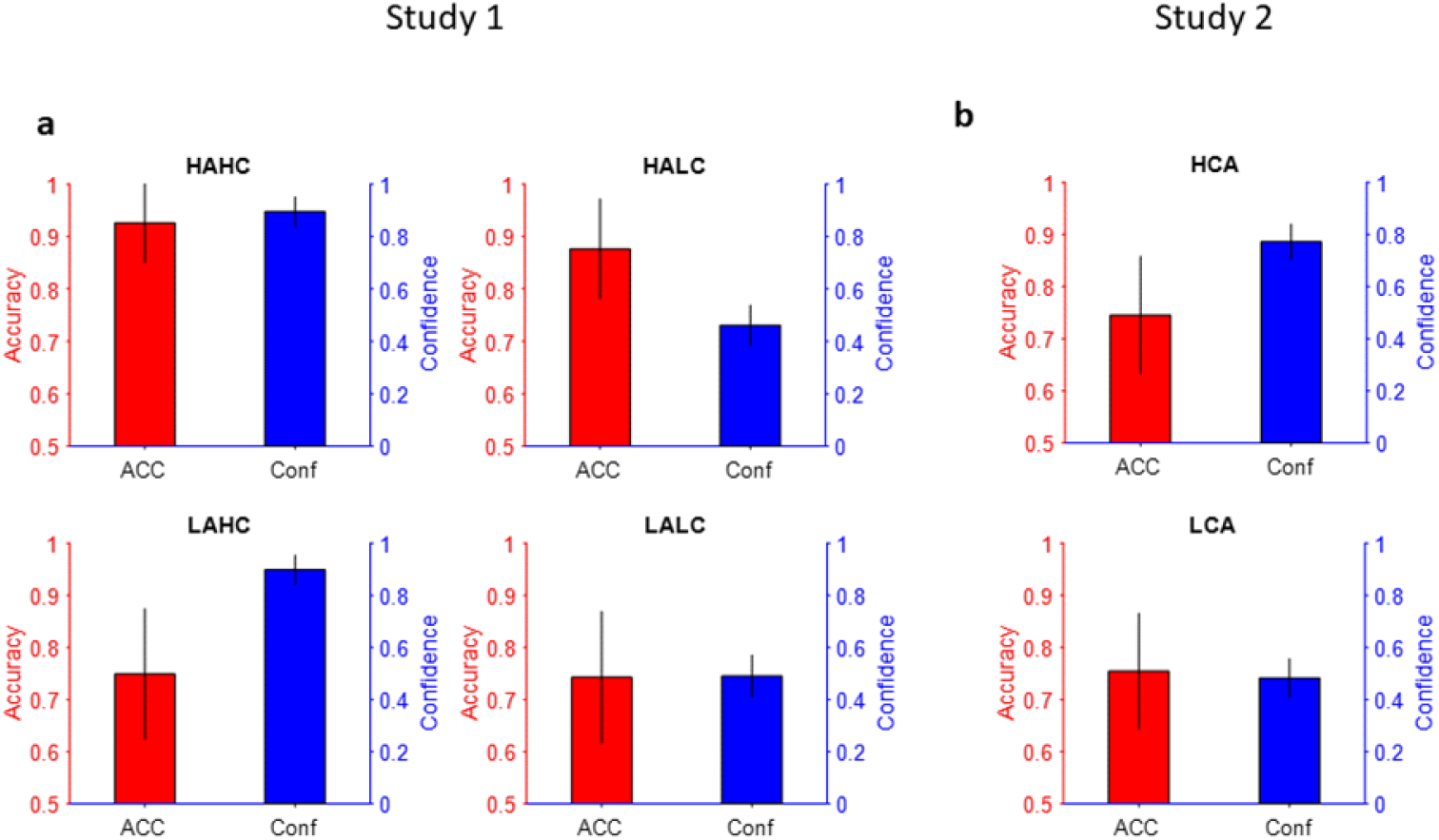
Accuracy and confidence of the Computer Generated Partners. Confidence is plotted in blue and accuracy is plotted in red. (a) Study 1 - HAHC: high accuracy and high confidence. HALC: high accuracy and low confidence. LAHC: low accuracy and high confidence. LALC: low accuracy and low confidence. Error bars: standard deviation across 12 CGPs. Top-right, Bottom-left and Bottom-right are same as Top-Left but for HALC, LAHC and LALC, respectively. (b) Study 2 - same as (a) but for 2 CGPs and across 15 partners generated for 15 participants. HCA: High confidence agent; LCA: low confidence agent

**Figure 1-figure supplement 2.**
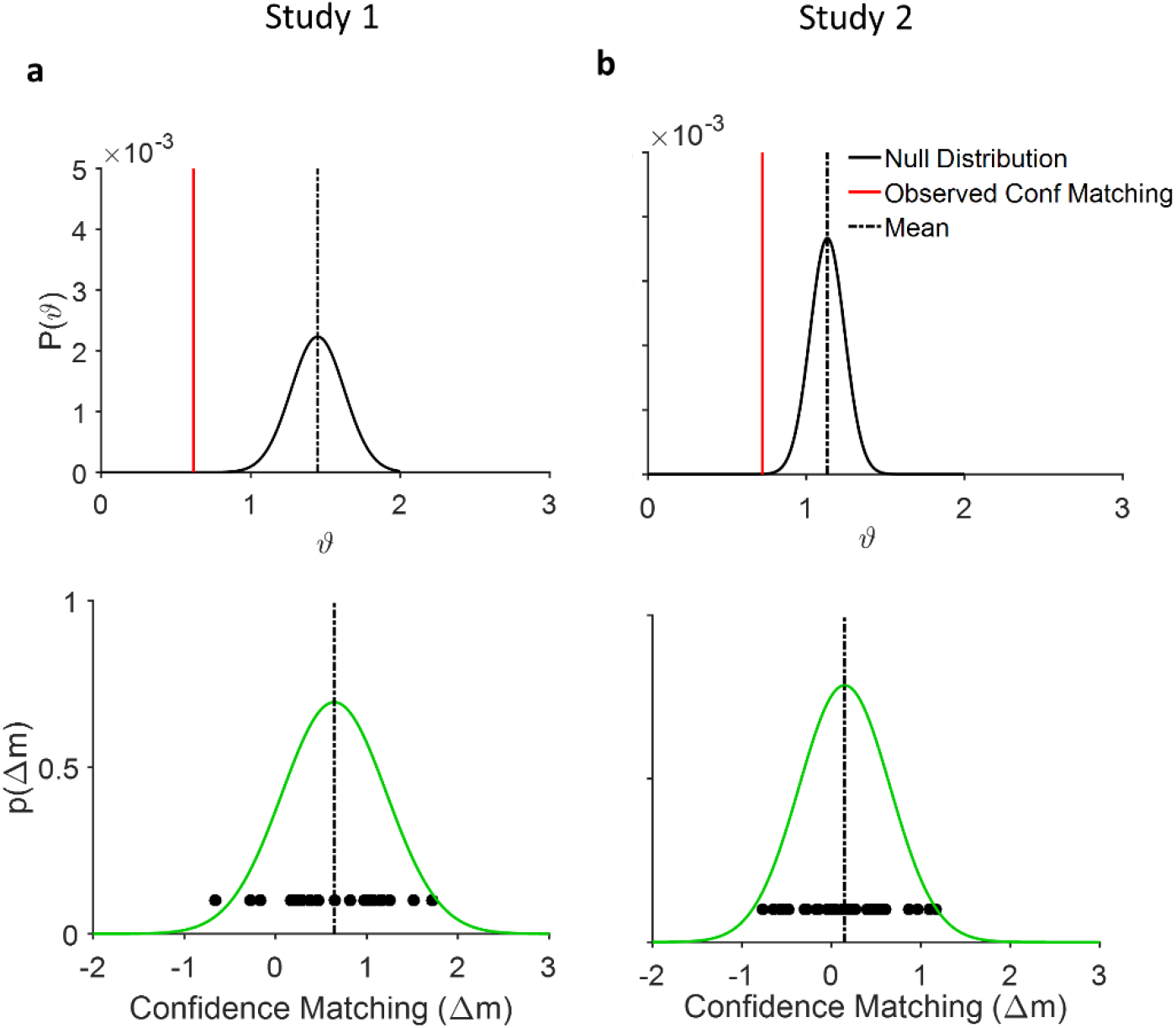
(a) Top: Permutation test. The empirically observed difference in mean confidence (red line) is significantly different from the distribution of the expected mean (black curve and dotted line) under null hypothesis (p(ϑ)), random pairing, (p∼0 for both studies). Bottom: Probability density function (green curve) over confidence matching index defined as (*Δm* = *C*_*isolated*(*Subject*)_ − *C*_*agent*_ − *C*_*social*(*Subject*)_ − *C*_*agent*_). Dots denote *Δm* for each combination of 12 participants by 4 CGPs. The mean confidence difference is significantly different from zero (study 1, p<0.001, paired t-test t(47)=5.5, study 2: p=0.11 t(29)=1.59 but significantly different for HCA condition, p<0.001 t(14)=4.02). (b) same as (a) but for the second study in which we had 2 CGPs

**Figure 1-figure supplement 3.**
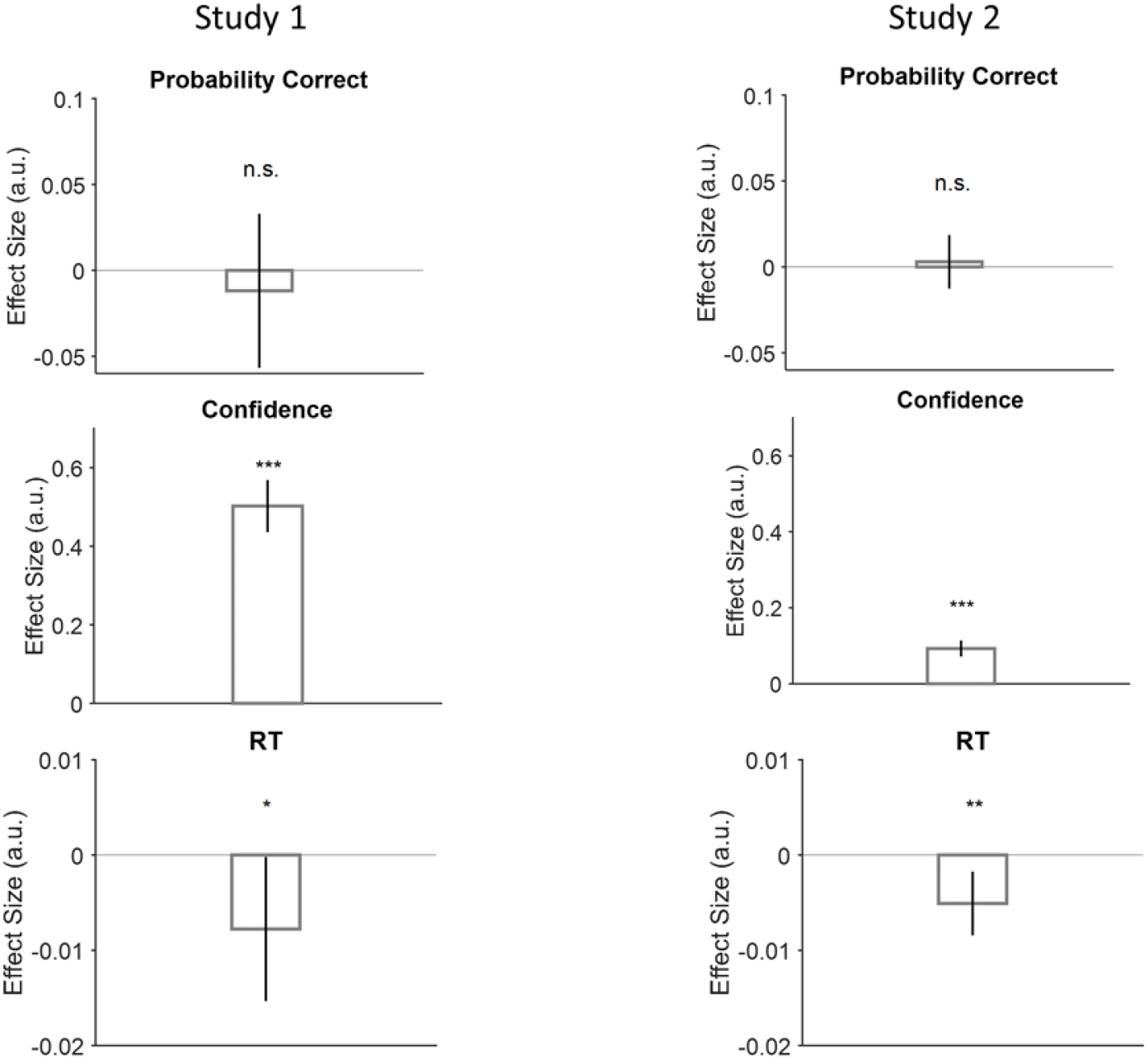
Direct examination of the hypothesis that the partner’s confidence at trial t modulates behaviors at trial t+1. (Probability correct: First row, Confidence: Second row and RT: last row) on study 1 (first column) and 2 (second column). We used a GLMM similar to figure 1.c and plotted the model’s coefficient regarding confidence of the previous trial (from 1 to 6). The results did not change compared to figure 1.c indicating the previous trials confidence impacts the behaviors in the upcoming trial regardless of experiment conditions (HCA, LCA)

**Figure 3-figure supplement 1.**
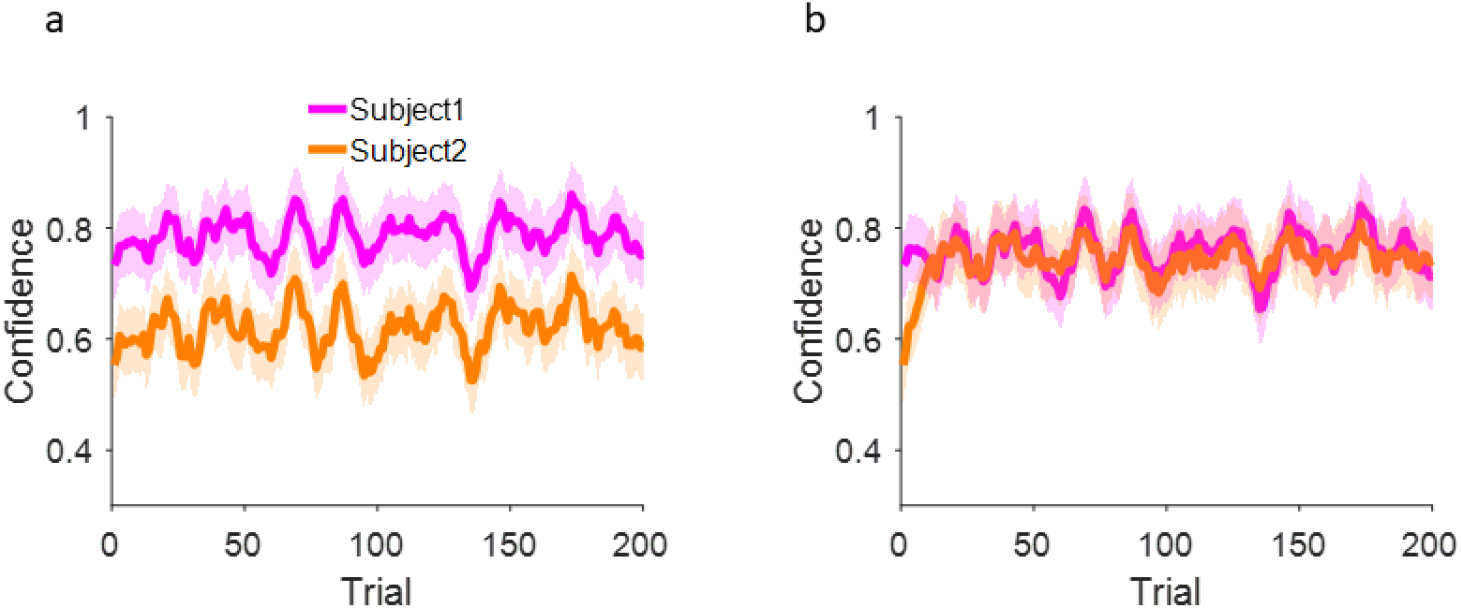
Confidence matching without removing the correlation with the shared stimulus coherence.

**Figure 3-figure supplement 2.**
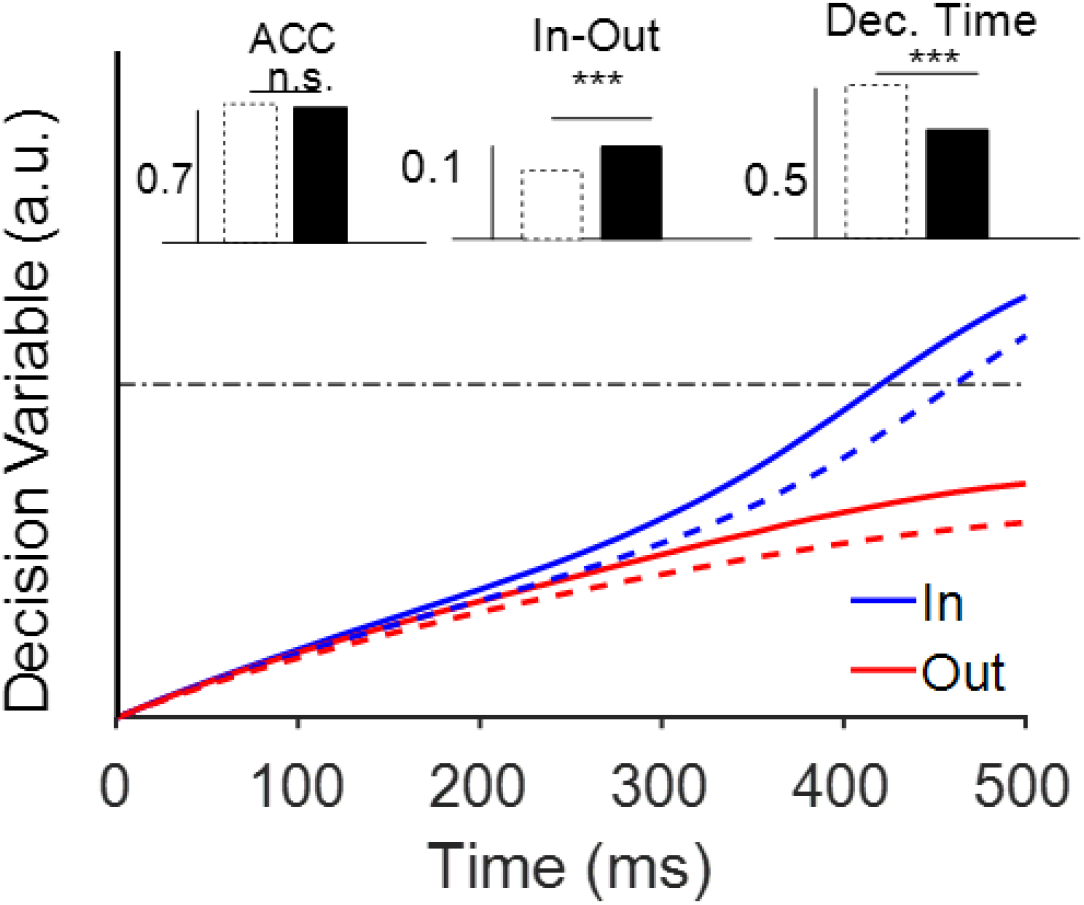
The effect of top-down current on the attractor network. The results of model simulations with a specific value of top down current (*W*_*x*_ = 0.003). This plot shows the average accumulated evidence of the model in 1,000 repetitions with (solid lines) and without (dashed lines) top-down current for a 0.5s duration of stimulus with 6.4% coherency. Importantly, the network was shut down after stimuli offset, receiving only the noise terms (equation 1 and 2) for 2 s (that is more than RTs of 99% of trials). The dot-dashed black line indicates the decision threshold. Inset plots show how the accuracy (ACC), absolute difference of firing rates of two populations (In-Out) and decision time (Dec. Time) changed in the presence (black bars) or absence (white bars) of the top-down current. (***) indicates p<0.001, paired t-test between runs; n.s. denotes that there is no significant difference between conditions.

**Figure 3-figure supplement 3:**
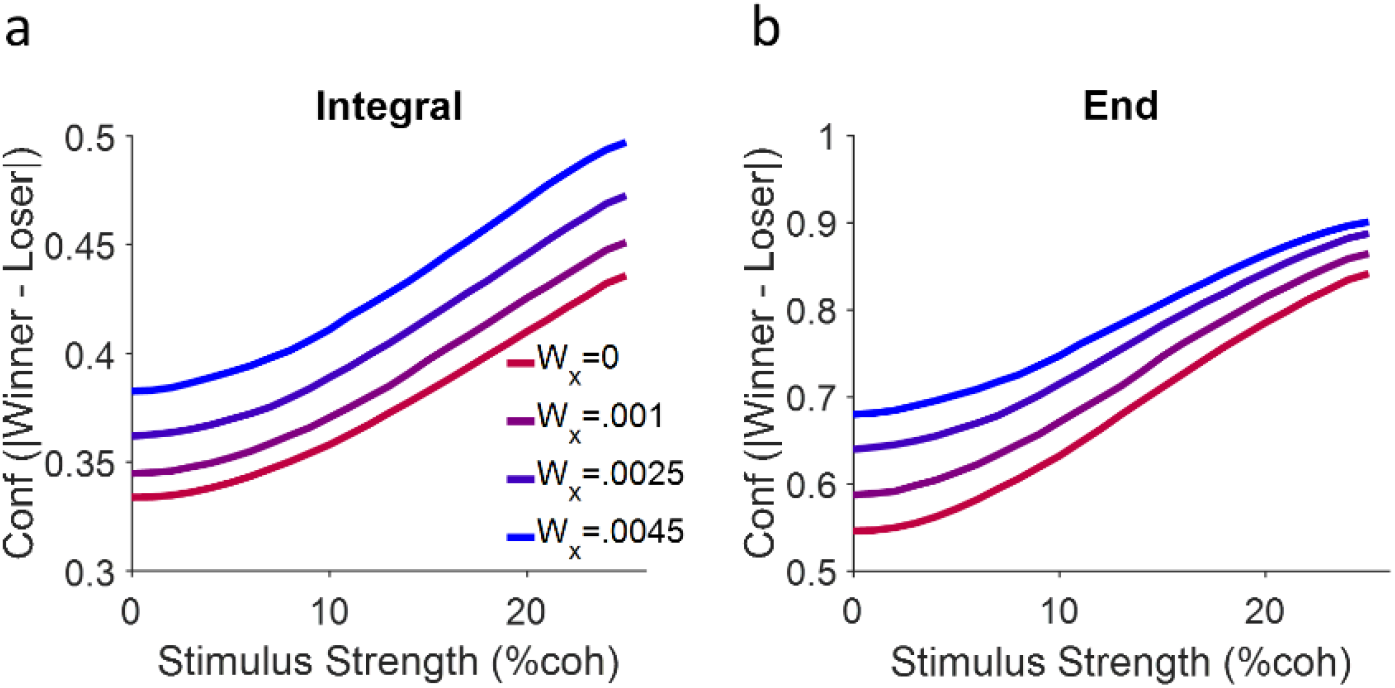
Model performance regarding different confidence representations. a) confidence representation based on equation 8 in the main text. b) same as a but here confidence is calculated as the absolute difference of the winner and loser signal but only at the end of the stimulus duration (500 ms). Each curve is a simulation 10,000 trials.

**Figure 3-figure supplement 4.**
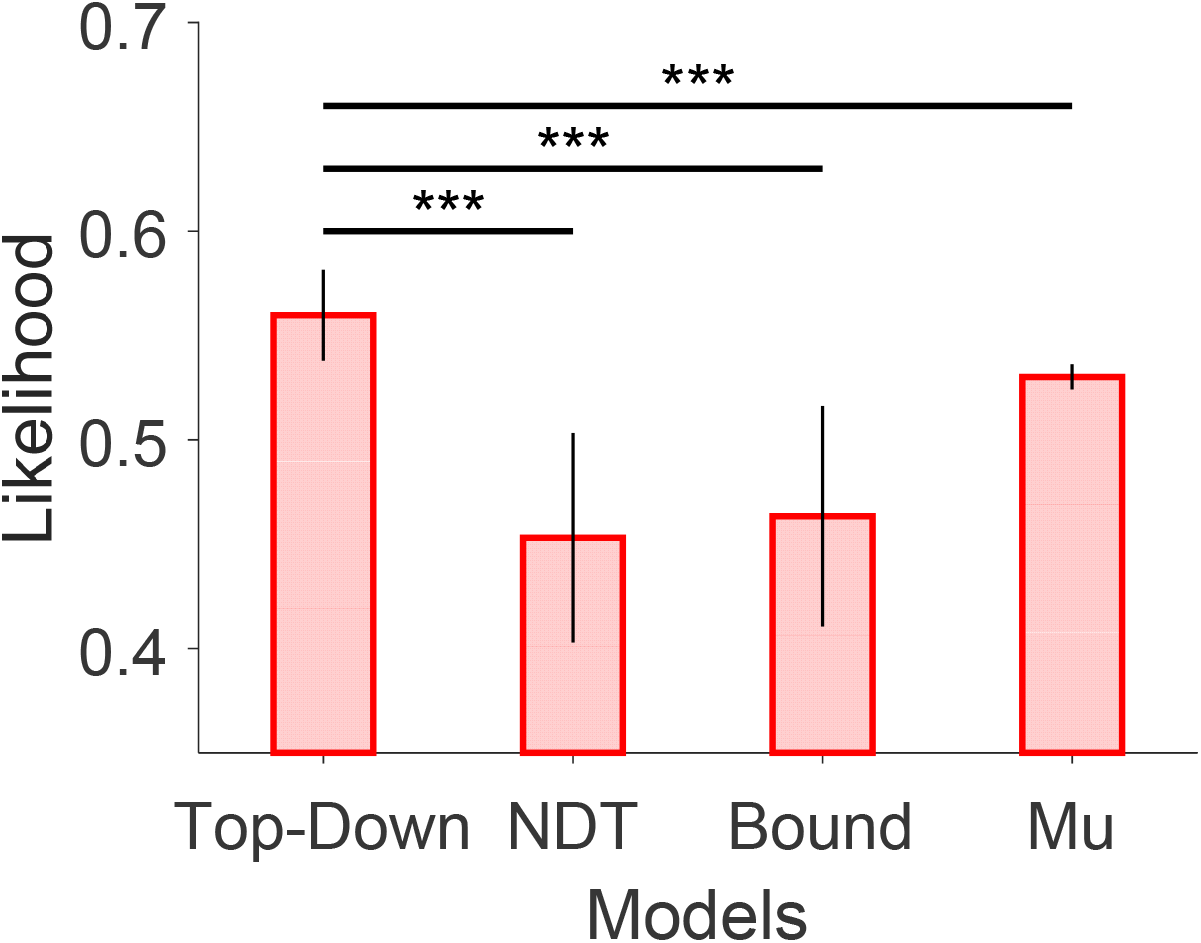
Comparison of likelihood of four models. Error bars are STD across 20 different initial points. (***) indicates p<0.001. (Wilcoxon Rank Sum test, n=20).

**Figure 3-figure supplement 5.**
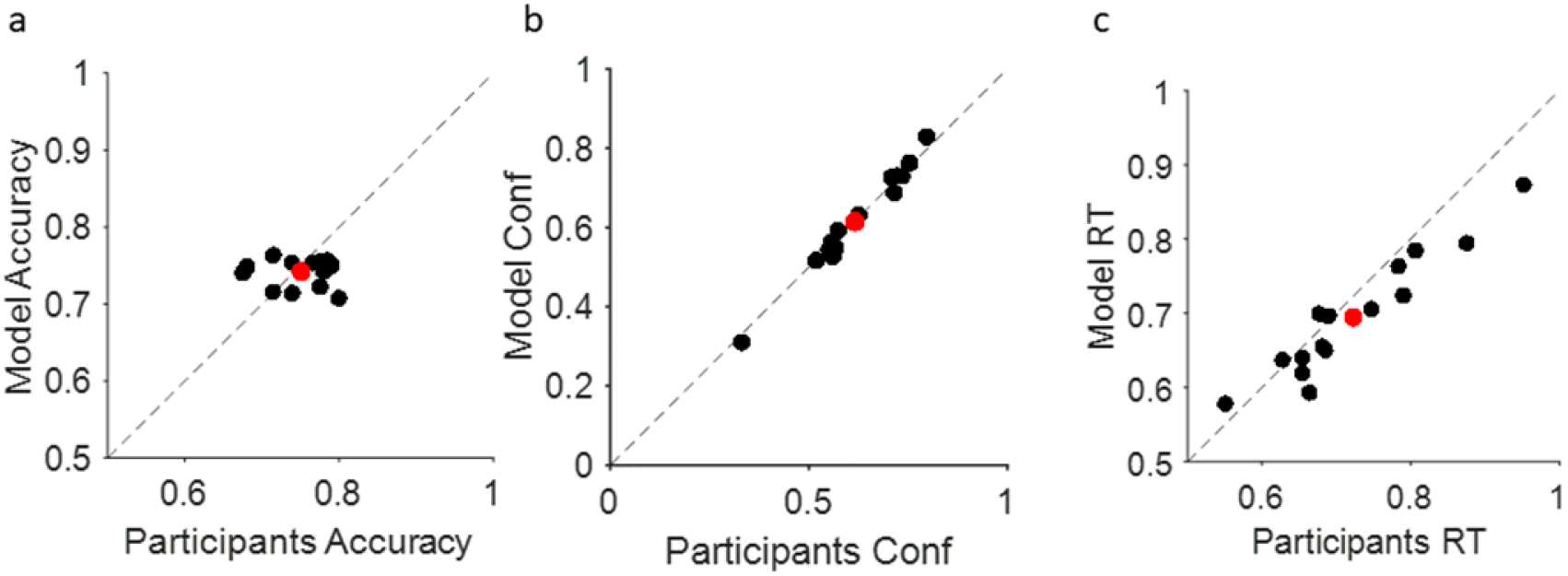
Comparison of the correspondence between the model fits (y-axis) with behavioral data (x-axis) of each participant (n=15). Accuracy (a), Confidence (b) and RT (c) in the Isolated session. Each dot represents a single participant. The red dot in each panel is the average. The model was able to capture behavioral trends across participants in which it showed no significant difference between model predictions and behavioral data (figure S12; p=0.13, 0.86, 0.40, for accuracy, confidence and RT respectively, Wilcoxon Ranksum Test n=15)

**Figure 4-figure supplement 1.**
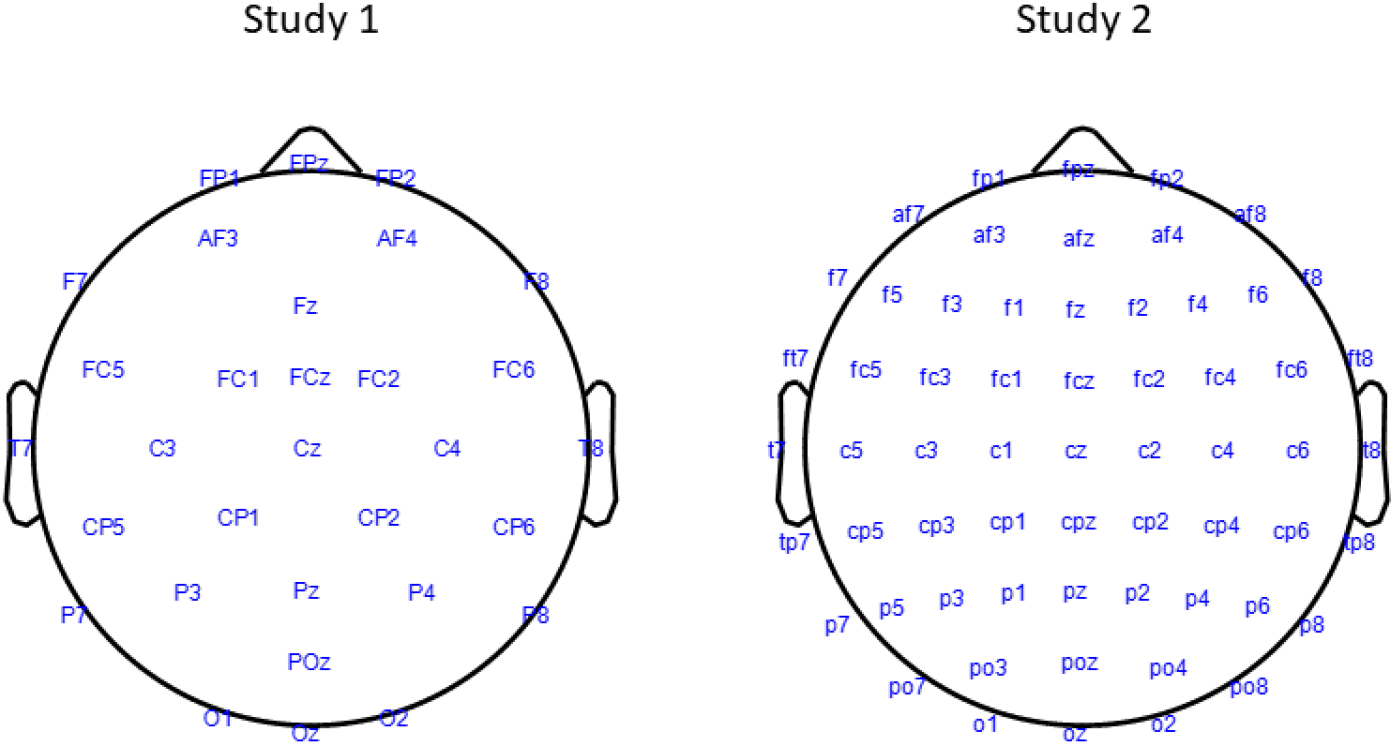
Electrode placement in each study.

**Figure 4-figure supplement 2.**
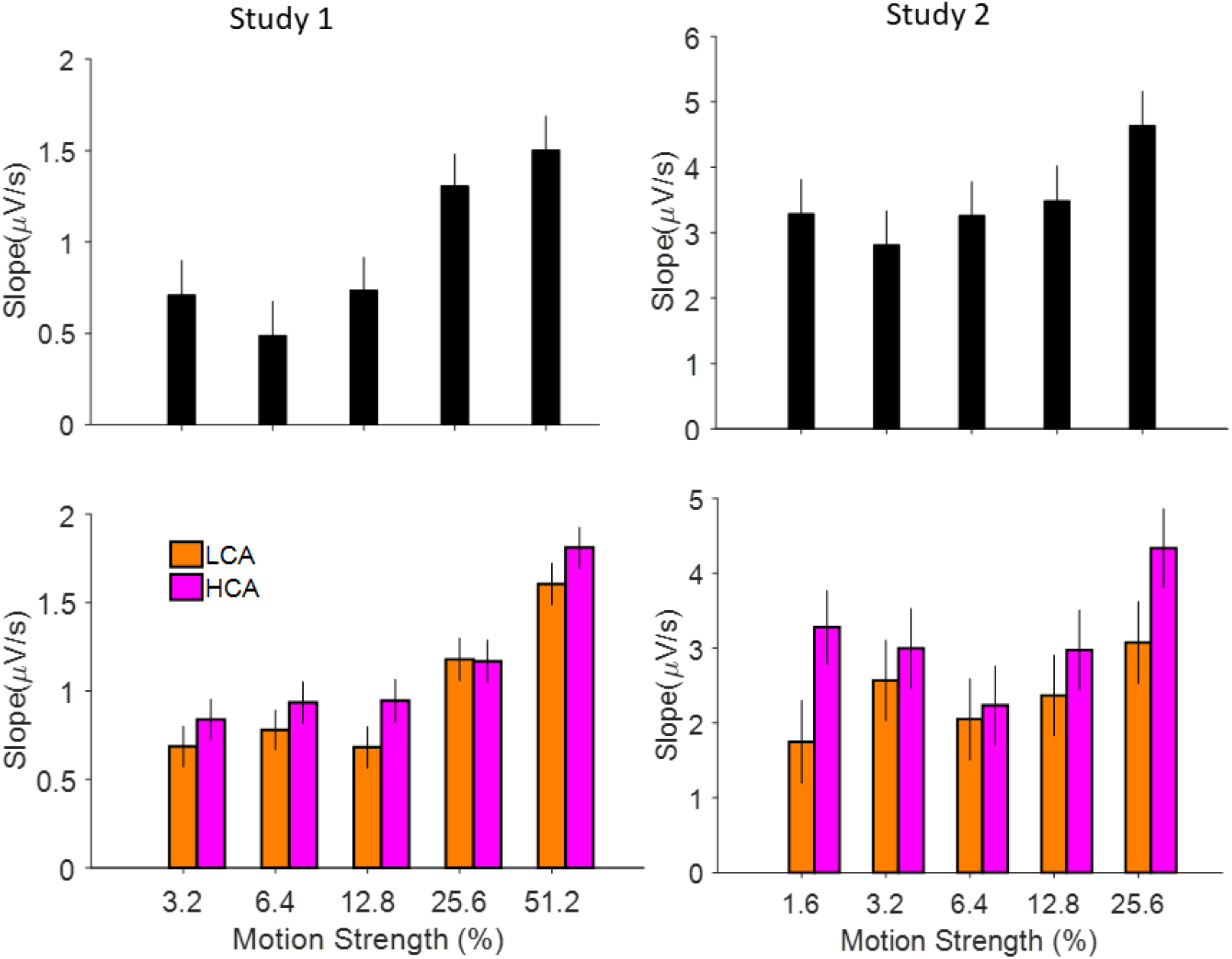
Relation of EEG signals from centro-partial area of the brain to coherence levels and social conditions. Top-Left, ramping activities of the signals (calculated by a linear regression of signals amplitudes and the time windows of 0-500 ms) is modulated by coherence levels (GLMM Study 1: p<0.001, Study 2 (Top-Right): p=0.01). Bottom Left: relation of signals slope and social condition (HCA vs LCA) based on different coherence levels. HCA shows a steeper slope (see figure 4 and table S3 for more details). Right column is the same as Left column but for study 2.

**Figure 4-figure supplement 3.**
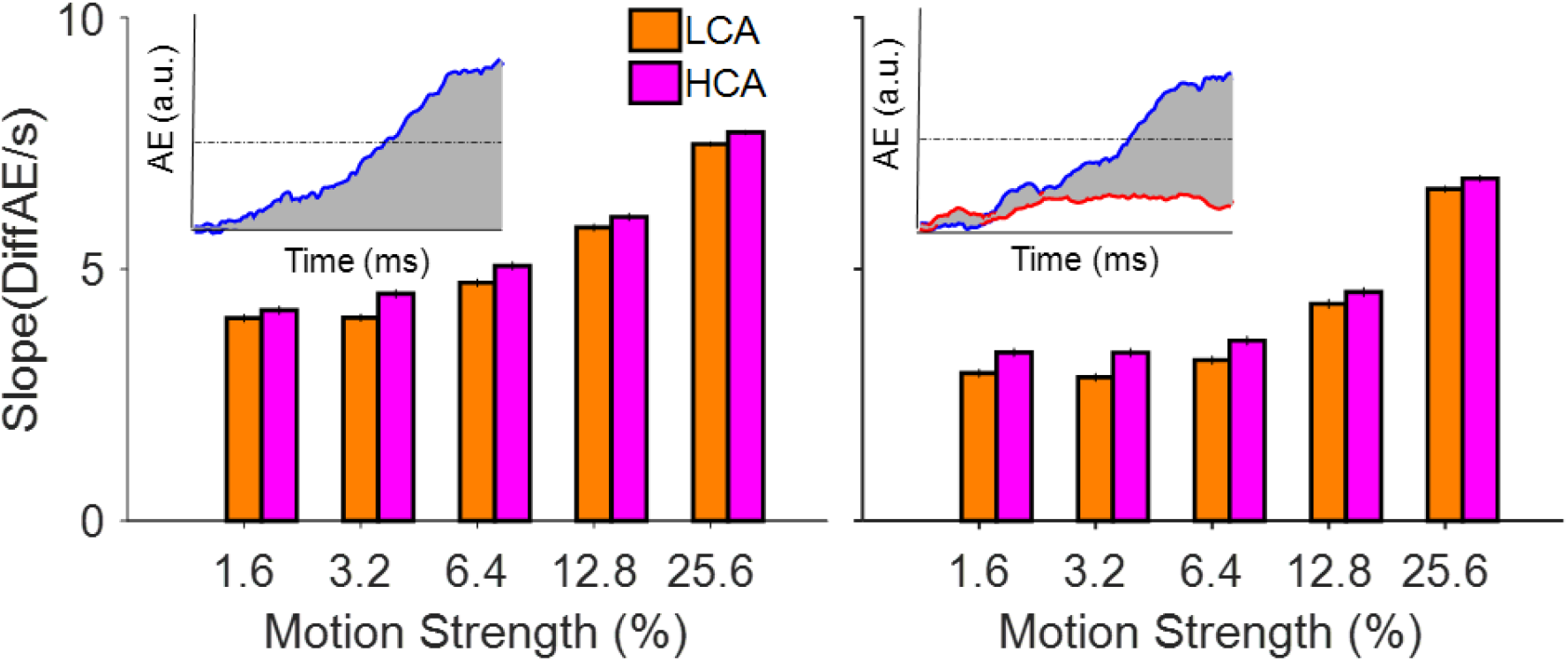
Simulated slope of the accumulator activity in our computational model in LCA and HCA conditions. (a) Slope of the winning accumulator (time window: 0-500 ms; shaded area, insets) at each coherence level for LCA and HCA condition. (b) Same as panel (a) but here for the *difference* in accumulator activities. Error bars are SEM across trials (n=3000, for each model).

**Figure 5-figure supplement 1.**
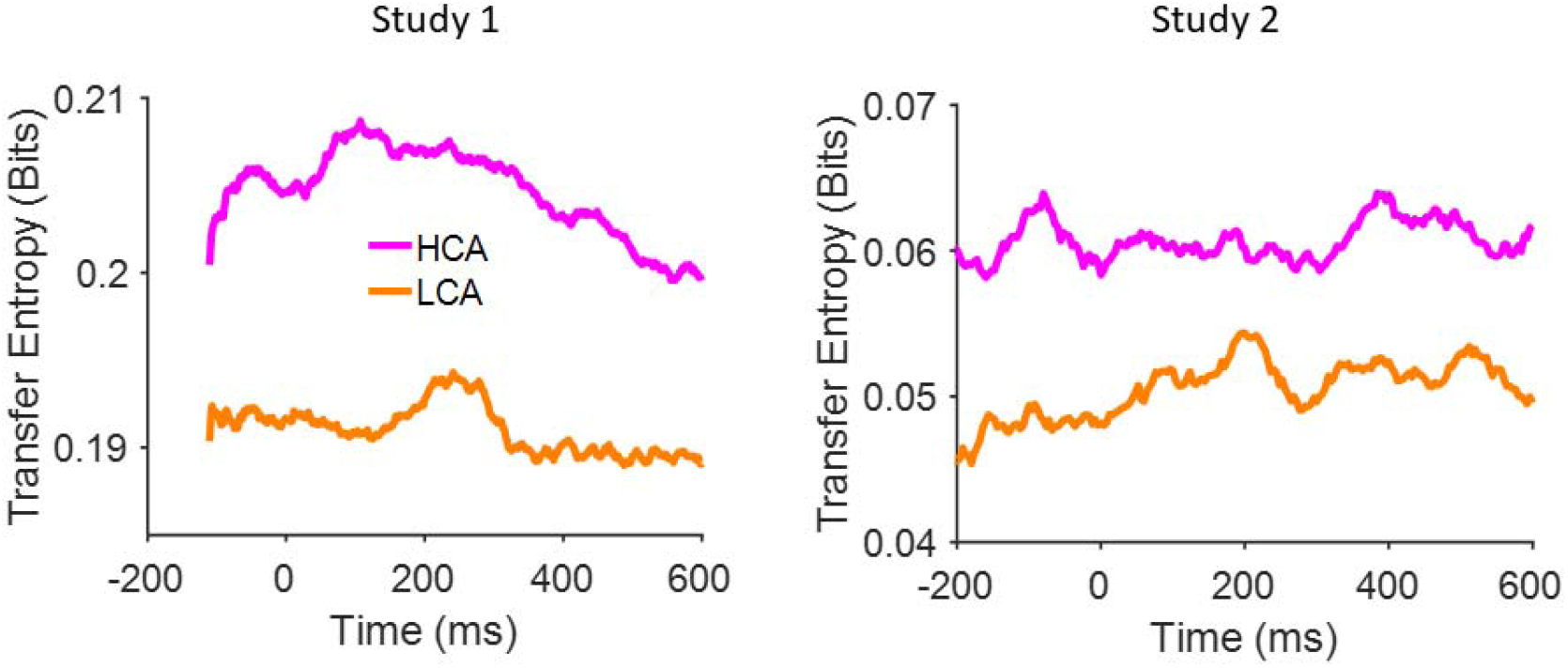
Temporal investigation of transfer entropy estimation under Social (HCA, LCA) conditions. HCA top-down connectivity is significantly higher than LCA (see figure 5 and table S4). Importantly, the transfer entropy estimation does not show any decisive relation to the time (x-axis) which is in line with our computational hypothesis of injecting a constant top-down current to the model.

